# Faulty Metabolism: A Potential Instigator of an Aggressive Phenotype in Cdk5-dependent Medullary Thyroid Carcinoma

**DOI:** 10.1101/2023.06.13.544755

**Authors:** Priyanka Gupta, Brendon Herring, Nilesh Kumar, Rahul Telange, Sandra S. Garcia-Buntley, Tessa W. Caceres, Simona Colantonio, Ford Williams, Pradeep Kurup, Angela M. Carter, Diana Lin, Herbert Chen, Bart Rose, Renata Jaskula-Sztul, Shahid Mukhtar, Sushanth Reddy, James A. Bibb

## Abstract

Mechanistic modeling of cancers such as Medullary Thyroid Carcinoma (MTC) to emulate patient-specific phenotypes is challenging. The discovery of potential diagnostic markers and druggable targets in MTC urgently requires clinically relevant animal models. Here we established orthotopic mouse models of MTC driven by aberrantly active Cdk5 using cell-specific promoters. Each of the two models elicits distinct growth differences that recapitulate the less or more aggressive forms of human tumors. The comparative mutational and transcriptomic landscape of tumors revealed significant alterations in mitotic cell cycle processes coupled with the slow-growing tumor phenotype. Conversely, perturbation in metabolic pathways emerged as critical for aggressive tumor growth. Moreover, an overlapping mutational profile was identified between mouse and human tumors. Gene prioritization revealed putative downstream effectors of Cdk5 which may contribute to the slow and aggressive growth in the mouse MTC models. In addition, Cdk5/p25 phosphorylation sites identified as biomarkers for Cdk5-driven neuroendocrine tumors (NETs) were detected in both slow and rapid onset models and were also histologically present in human MTC. Thus, this study directly relates mouse and human MTC models and uncovers vulnerable pathways potentially responsible for differential tumor growth rates. Functional validation of our findings may lead to better prediction of patient-specific personalized combinational therapies.

**Graphical abstract:** 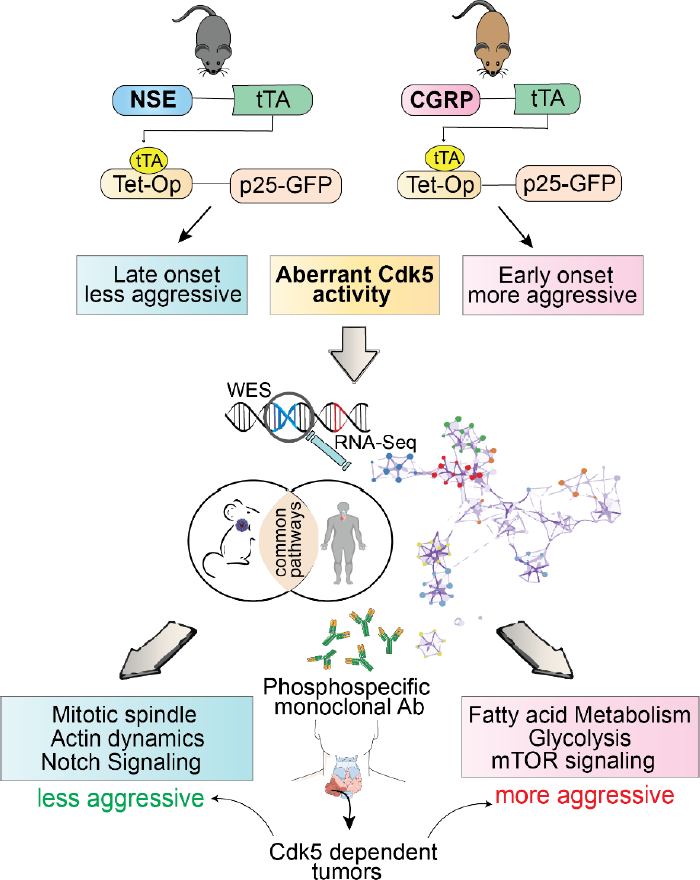

**Highlights:**

- CGRP driven aberrant Cdk5 activation develops early onset aggressive MTC
- Genetic alterations in mouse and human tumors disrupt common pathways
- Aggressive tumor model characterized by alterations in metabolic pathways
- Slow growing tumor model elicits disruption of mitotic spindle assembly

## Introduction

Medullary thyroid carcinoma (MTC) is derived from calcitonin-secreting parafollicular neuroendocrine (NE) cells. These tumors occur as sporadic or hereditary forms with an incidence rate of approximately 75% and 25%, respectively. MTC patients present clinically heterogeneous disease courses ranging from indolent to highly aggressive. The survival rate of 10-years varies from 100% (stage I) to 21% (stage IV) [1]. MTC accounts for 5-10% of all thyroid malignancies and is frequently associated with germline mutations in the *RET* proto-oncogene. Patients may also harbor somatic mutations in *HRAS, KRAS*, or *NRAS* [2]. The 5-year survival rate is ∼40% in patients with metastatic disease where tumors can spread to the cervical lymph nodes, or distant sites such as bones, lungs, liver, and brain [3–5]. Surgery is the only curative therapy for MTC, but resection of isolated metastases or other newer treatments have shown promise [6, 7]. However, a proper therapeutic regimen for aggressive, recurrent, and metastatic disease is still in abeyance.

RET mutations, serum calcitonin, and carcinoembryonic antigen (CEA) are known prognostic markers for MTCs. However, risk stratification based on serum biomarkers, namely calcitonin, CEA, carbohydrate antigen 19.9, or Ki67 expression has proven inefficient in identifying aggressive phenotypes, patients requiring immediate treatment, or resistance to existing therapeutic modalities [8]. The continuous global increase in MTC incidence corresponds with spiking mortality rates. Furthermore, epidemiological evidence from the past three decades has not indicated improvement in MTC diagnosis or overall patient survival. Lack of adequate predictive biomarkers, inconsistent long-term prognostic factors, and poor identification of aggressive phenotype attributed to the indolent course of the disease. Likewise, the unpredictable clinical behavior of patients triggers the overarching need for reliable biomarkers to detect aggressive forms of tumors.

Cyclin-dependent kinase 5 (Cdk5), and its co-activators p35/25 have emerged as critical molecular players in the tumorigenesis [9]. We have shown that aberrant activation of Cdk5 under the control of NE cell-specific promoter develops clinically accurate neuroendocrine tumors (NETs) [10–12]. In this study, we established a new Cdk5-driven model that exhibits early tumor onset and increased tumor volume doubling time compared to the previously established model. Our model replicates the aggressive phenotype observed in humans. A comprehensive approach deploying animal models, human tumors, and multi-omic analysis may lead to the identification of critical molecular candidates that can be utilized as biomarkers for diagnostics or potential targets for therapeutic intervention.

## Material and methods

### Generation of transgenic MTC models

The NSE-p25 bitransgenic mice were generated as described previously [13]. PiggyBac technology (Cyagen Biosciences) was used to generate single-copy CGRP transgenic mice in C57BL/6 background. Bitransgenic mice were then generated by crossing the TetOp-p25GFP mouse with that of CGRP-tTA or NSE-tTA. p25OE was controlled by doxycycline administration (Dox, 0.1 g/L) dissolved in drinking water. Doxycycline was removed at three weeks of age inducing tumors to grow for ∼10 weeks. Tumors were arrested by re-administration of doxycycline. At the end of the experiments, bilateral tumors were harvested and snap-frozen for sequencing experiments, immunoblotting, and fixed for immunohistological staining. All mice were group-housed on a 12 h light/dark cycle with access to food and water *ad libitum*. All animal procedures were performed under protocols approved by the UAB Institutional Animal Care and Use Committee.

### Polymerase Chain Reaction

Positive pups carrying CGRP-tTA transgene confirmed by genotyping using primers-forward (F): ATCAAGAGTCACCGCCTCGC; Reverse (R): TTTGAGCGAGTTTCCTTGTCGTC. Transgene product size 215 bp. All positive pups were confirmed by PCR to not contain any integration of the helper plasmid. The pups carrying the NSE-tTA transgene were confirmed using the following primers – tTA 1080R: TTT CTG TAG GCC GTG TAC CTA; tTA 906F: GAT GTT AGA TAG GCG CCC TAC TCA C; Gdf5-D1: GGA GCA CTT CCA CTA TGG GAC & Gdf5-D2: AAA GAG TGA GGA GTT TGG GGA G. Transgene product size 243 bp. The Tet-op p25 gene was evaluated by the following primers – HS 18: CCA TCG ATC TAG TAC AGC TCG TCC ATG C; HS 28: AAG GAC GAC GGC AAC TAC; Gdf5-D1: GGA GCA CTT CCA CTA TGG GAC & Gdf5-D2: AAA GAG TGA GGA GTT TGG GGA G. Transgene product size 400 bp. Bitransgenic mice were positive for both the NSE or CGRP and p25-GFP alleles while control littermates were positive only for one of the two alleles. All reactions were carried out using the 2X master mix from Promega.

### Magnetic Resonance Imaging

MRI was performed with a Bruker Biospec 9.4 Tesla instrument using Paravision 5.1 software (Bruker Biospin, Billerica, MA). A Bruker 72 mm ID volume coil was used for excitation and a custom 24 mm surface coil for signal reception (Doty Scientific Inc., Columbia, SC). Mice were anesthetized with isoflurane gas and respiration observed with a MRI-compatible physiological monitoring system (SA Instruments Inc., Stony Brook, NY). Animals were imaged in supine position on a Bruker animal bed system with circulating heated water to maintain body temperature. A 2D T2-weighted RARE sequence was used for imaging of the abdomen. The following imaging parameters were used: TR/TE = 2000/25ms, echo spacing = 12.5ms, ETL = 4, 2 averages, 29 contiguous axial slices with 1 mm thickness, FOV = 30x30 mm and matrix = 300x300 for an in-plane resolution of 100 µm. Prospective respiratory gating was used to minimize motion artifacts. Tumor volumes were quantitated using ImageJ software.

### Immunoblotting and immunohistological staining

Cells and tumor tissues were lysed in 1% SDS plus 50 mM NaF. Samples were sonicated briefly, spun at 20,000 g for 5 min, and supernatant combined with Laemmli buffer for analysis by SDS-PAGE followed by transfer to nitrocellulose membrane and subsequent detection of target proteins using a Li-Cor Odyssey imaging system. Immunoblotting was performed using antibodies for Cdk5 (Rockland 200-301-163; RRID:AB_11182476; 1:1000), GFP (Cell Signaling Technology 2956; RRID:AB_1196615; 1:2000), P-T202 LARP6 (Bibb Lab; 1:1000), LARP6 (Invitrogen PA5-41881; RRID:AB_2605747; 1:1000), P-S17 H1.5 (Bibb Lab; 1:1000), H1.5 (Santa Cruz sc-247158; RRID:AB_10847577; 1:1000), P-S988 RBL1 (Bibb Lab; 1:1000), RBL1 (Santa Cruz sc-318; RRID:AB_2175428; 1:500), P-S391 SUV39H1 (Bibb Lab; 1:1000), SUV39H1 (sc-377112; RRID N/A; 1:1000), P-T143 FAM53C (Bibb Lab; 1:1000), FAM53C (Invitrogen PA5-114093; RRID:AB_2884608; 1:1000) and β-actin (Invitrogen AM4302; RRID:AB_437394; 1:5000).

For immunostaining, tissues were fixed in formalin, embedded in paraffin, and sliced into 5 µm sections for placement on glass slides. Samples were deparaffinized and subjected to high temperature antigen retrieval in citrate buffer (pH 6.0). For IHC, samples were permeabilized in 0.3% Triton X-100, blocked with 5% normal goat serum, and then incubated overnight at 4°C in primary antibodies to GFP (Cell Signaling 2956; 1:200), ChrA (Abcam ab15160; 1:1000), P-T202 LARP6 (Bibb Lab; 1:200), P-S17 H1.5 (Bibb Lab; 1:50), and P-S988 RBL1 (Bibb Lab; 1:100) diluted in 5% normal goat serum and 0.3% Tween 20. Sections were then incubated in 0.3% hydrogen peroxide and biotinylated secondary antibodies (Pierce 31820 or 31800; 1:500) applied to slides for 1 h at room temperature followed by 30 min of streptavidin-HRP. Slides were then incubated with DAB Chromogen (Dako Liquid DAB+ substrate K3468) and counter-stained with hematoxylin. Standard procedures were used for H&E staining (Feldman and Wolfe, 2014). Archival human tissues and tissue microarrays were obtained in accordance with UAB IRB protocol IRB-300002147. Stains of human tissues were reviewed for quality by a board-certified pathologist with expertise in thyroid pathology.

### Whole exome sequencing (WES) and RNA sequencing (RNA-Seq)

Total DNA was extracted from the frozen tumors using DNeasy Blood and tissue kit (Qiagen) according to the manufacturer’s instructions, and RNA was purified using RNEasy Plus Mini Kit (Qiagen). For WES, exome capture was performed using the Agilent SureSelect Mouse All Exome QXT capture kit (Agilent). Briefly, the genomic DNA was subjected to tagmentation reactions inserting adaptor sequences randomly throughout the genome. The DNA was PCR amplified and then incubated with biotin labeled RNA capture probes complementary to every exon. Following purification of the exome sequences through streptavidin-magnetic bead separation, the DNA was amplified with primers that introduced 8-nucleotide index so that separate samples can run in the same lane for sequence analysis. The exomic libraries were run on the NextSeq500 next generation sequencer from Illumina (Illumina, San Diego, CA) with paired end 75 bp reads using standard techniques. RNA-seq was performed on the same instrument. Briefly, RNA quality was assessed using the Agilent 2100 Bioanalyzer. RNA Integrity Number (RIN) of ≥ 7.0 was used for sequencing library preparation. Quality controlled RNA was converted to a sequencing ready library using the NEBNext Ultra II Directional RNA library kit with polyA selection as per the manufacturer’s instructions (New England Biolabs). The cDNA libraries were quantitated using qPCR in a Roche LightCycler 480 with the Kapa Biosystems kit for Illumina library quantitation (Kapa Biosystems, Woburn, MA) before cluster generation.

### Bioinformatics analysis

<u>Exome Seq</u>– MoCaSeq pipeline was used to analyze raw WES data (source code: https://github.com/roland-rad-lab/MoCaSeq)[14]. Using Docker and Ubuntu Linux, the pipeline was set up. With Trimmomatic (v0.38) [15] and BWA-MEM (v0.7.17)[16], the raw reads were aligned to the mouse reference genome GRCm38.p6. For further post-processing, Picard 2.20.0 and GATK (v4.1.0.0) were used [17]. The cutoff of the variant allele frequency was set at ≥10% as recommended [14]. For the loss of heterozygosity (LOH) analyses from WES data, somatic SNP calling was performed using Mutect2[18]. LOH analyses were limited to reads with a mapping quality of 60 to avoid ambiguous SNP positions caused by mis-mapping. CopywriteR (v2.6.1.216) [19], which extracts DNA copy number information from off-target reads, was used to call CNVs. Custom Python (v.3.10) and Shell scripts were used for downstream analysis and visualization. <u>RNA Seq</u>– To remove low-quality reads from raw sequences, fastp (v0.21.0) was used [20]. Sequence alignment was performed using STAR v2.7.3a-GCC-6.4.0-2.28 aligner[21] and GRCm39 assembly. Using the accepted alignment hits, gene counts were obtained using HTSeq (HTSeq v0.12.3-foss-2018b-Python-3.6.6)[22]. The differentially expressed genes (1.5 < = Fold change and 0.05 = > FDR) were identified using DESeq2 (DESeq2_1.36.0) [23] and R (v4.2.0). The enriched gene sets and pathways were analyzed using GSEApy (0.13.0), Enrichr, Shiny GO, and Metascape.

### Cell proliferation

Cell proliferation assay was performed on mouse MTC cells[10] in the presence or absence of doxycycline using CyQUANT™ Direct Cell Proliferation Assay following the manufacturer’s protocol (Thermo Fisher Scientific)[24].

### Generation and purification of monoclonal antibodies

The rabbit monoclonal antibodies were developed directly from isolated B cells of immunized animals without the use of hybridomas. Briefly, an antigen peptide containing the phosphorylated site of interest is synthesized with an N-terminal cysteine and conjugated via the thiol-group to carrier proteins. At least two rabbits are immunized with the peptide. After immunization and subsequent boosts, peripheral blood is drawn and the titer of the antiserum against the antigen is determined via indirect ELISA assays against the phosphorylated peptide. The rabbit with the highest titer and desired activities was used for the isolation of peripheral blood mononuclear cells (PBMCs). Antigen-specific B cells are cultured *in vitro* in multi-well plates and supernatant samples are screened to identify desired antibodies. The cDNAs encoding the heavy (H) and light (L) chains of the antibodies were obtained by reverse transcription-polymerase chain reaction (RT-PCR) of RNA samples isolated from positive B cell clones. H and L cDNAs cloned into mammalian expression vectors and were transiently transfected into Chinese hamster ovary (CHO) cells. Recombinant monoclonal antibodies were generated from the expressed heavy and light chains. Before purification, recombinant antibodies were screened by ELISA and additional application assays. For antibodies that are specific for phosphorylated sites on proteins, the non-phosphorylated antigens were used for counter-screening assays (monoclonal rabbit antibodies by Excel BioPharm LLC). The antibody clones have been submitted to CPTAC Antibody Portal (National Cancer Institute Clinical Proteomic Tumor Analysis Consortium) designated as #CPTC-FAM53C-1, #CPTC-SUV39H1-1, #CPTC-H1-5-1, #CPTC-LARP6-1, and #CPTC-RBL1-1.

### Statistics

Statistical analysis was performed using Prism 8.4.2 (GraphPad Software). Student’s *t*-test was used to determine the significant difference between the two groups. Experimental replicates or sample sizes presented as n are provided in the figure legends. Differences between the groups were considered significant at p < 0.05. The degrees of significance were reported as *p < 0.05, **p < 0.01, ***p < 0.001.

## Results

### CGRP promoter driven aberrant Cdk5 develops aggressive tumors

While Cdk5 is widely implicated in brain disorders, its tumorigenic function is still nascent. To gain insights into the functional role of Cdk5 in tumor progression at the genomic and transcriptomic level, we engineered bi-transgenic mouse tumor models where tumorigenesis is induced by aberrant Cdk5 activation. A doxycycline-regulated system was deployed where NE cell-specific promoter-driven tetracycline transactivator (tTA) induces p25 overexpression (p25OE) resulting in tumor development at the orthotopic site. Eventually, p25OE triggers reciprocity of Cdk5 with p25 over p35 rendering aberrant kinase activity. The resultant Cdk5/p25 interaction facilitates pro-neoplastic signaling compared to its physiological counterpart, *i.e.* Cdk5/p35 (Figure 1A).

**Figure 1.**
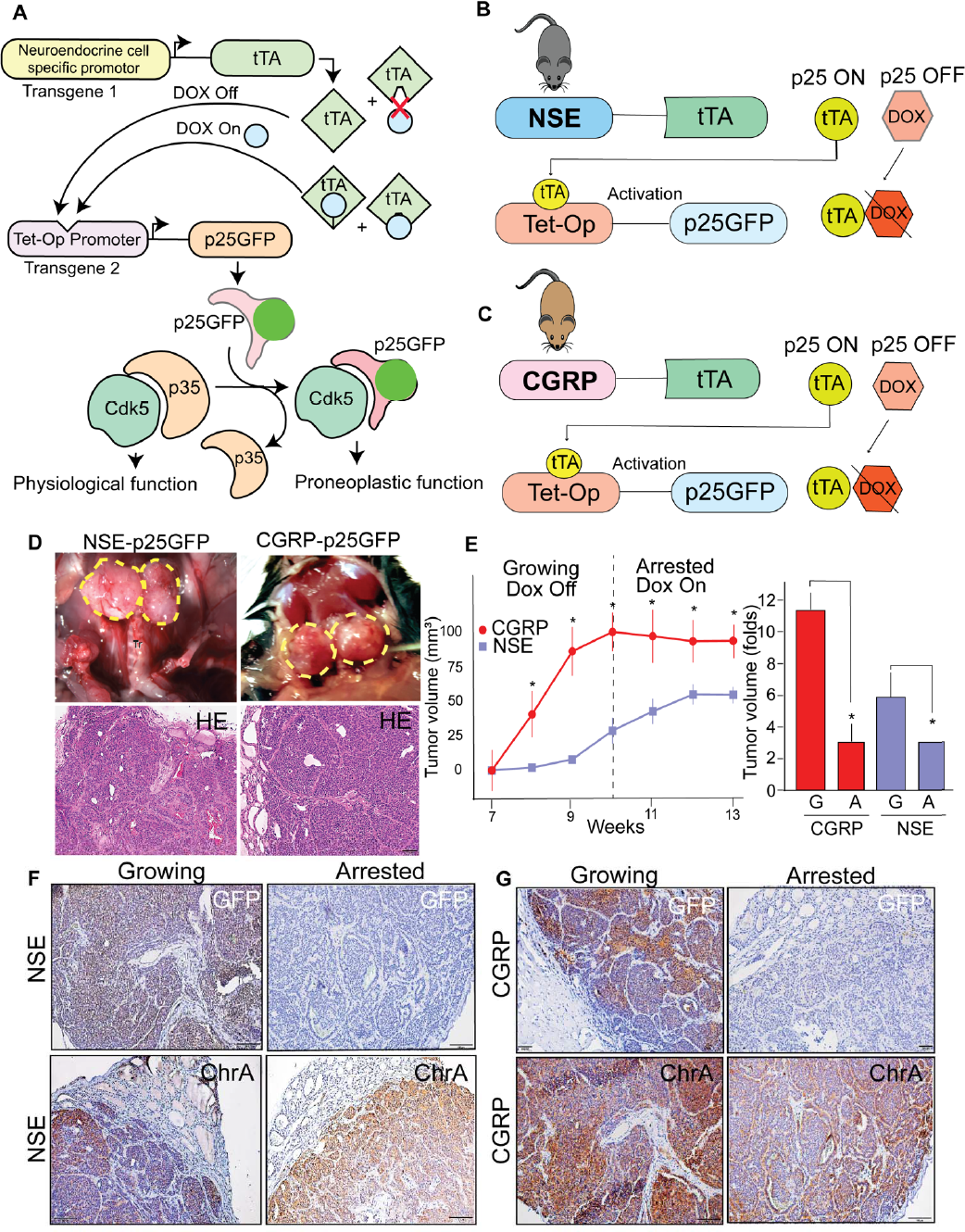
Generation of transgenic mouse models of MTC. (A) Schematic showing tetracycline controlled bitransgenic system where a neuroendocrine cell-specific promoter linked to the tetracycline transactivator (tTA) activates Tet-Operon driving p25-GFP expression. The resultant Cdk5–p25GFP interaction promotes neoplastic transformation. (B-C). Schematic showing the induction of p25GFP in doxycycline-controlled bitransgenic models driven by NSE and CGRP promoters. (D) Gross anatomy of tumors in NSE-p25GFP and CGRP-p25GFP models (upper panel); histopathological neuroendocrine features characterized by H & E (bottom panel), scale; 50μm, tr; trachea. (E) Tumor volume growth curves of NSE (growing n=4; arrested n=5) and CGRP (growing n=3; arrested n=5) mice showing weekly measurements of tumor volume under Dox Off vs. Dox On conditions (left); Bar graph on right shows tumor volume fold change of growing tumors (G; 14 weeks Dox Off) relative to arrested (A; 14 weeks Dox On); values are mean ± SD, *p<0.05 compared by Student’s *t*-test. (F-G) Representative immunostains comparing expression of p25GFP and Chromogranin A (ChrA) in NSE (F) and CGRP (G) mice tissues extracted from growing and arrested tumors. A version of NSE-p25GFP MTC tumors (1D, left) was previously published [10] and shown here at a different magnification for comparison.

We previously showed that p25OE in calcitonin-secreting C cells controlled by the neuron-specific enolase (NSE) promoter developed bilateral MTCs in mice [10] (Figure 1B). Calcitonin gene-related peptide (CGRP) is a splice variant of the calcitonin gene which translates into a neuropeptide localized in neuronal and neuroendocrine cells. Studies have shown that CGRP promoters are more efficient in restricting the transgene expression in calcitonin-secreting C cells in comparison to neuronal cells [25, 26]. Hence, we exploited the CGRP promoter in our transgenic system to induce stringent p25OE in thyroid C cells while preventing leaky expression in off-target tissues (Figure 1C). Indeed, both NSE and CGRP promoter-driven p25OE developed MTC tumors (Figure 1D). However, the tumor onset and growth rate of CGRP-p25OE model was significantly higher than that of NSE-p25OE. The tumor size in CGRP was ∼100mm^3^ vs. 25mm^3^ in NSE at 10 weeks after doxycycline removal, showing a ∼4-fold tumor growth rate increase in CGRP-p25OE mice (Figure 1E). Both murine models overexpressed p25GFP and chromogranin A (Chr A) in growing tumors signify typical NETs features (Figure 1F-G). Of note, p25GFP expression was drastically decreased in arrested conditions confirming the dependency of tumor growth on Cdk5 (Figure 1F-G). The comparable p25GFP expression between NSE and CGRP growing tumors suggests the involvement of other factors causing rapid tumor growth in CGRP over the NSE model. Our data show that CGRP-p25OE develops early-onset aggressive forms of MTCs compared to the previously reported NSE-p25OE mice. These results prompted us to examine the downstream nodes of Cdk5 signaling in these models to identify plausible effectors associated with rapid tumor growth.

### Genomic alteration landscape in mouse and human tumors

We performed whole-exome sequencing (WES) to determine the mutational profile of NSE and CGRP MTC models. Both models displayed highly heterogeneous mutational profiles across the chromosomes (Figure 2A and Table S1) where NSE-p25OE harbored ∼876 somatic mutations compared to ∼50 in CGRP-p25OE (Figure 2B). Of these, the majority include 154 missense, 176 silent, and 66 5’UTR mutations in NSE-p25OE. In contrast, CGRP-p25OE accumulated 12 missense, 9 silent, and 6 5’UTR (Figure 2B). Furthermore, exome data displayed drastic differences in the number of SNPs, insertions, and deletions between NSE and CGRP tumors, likely causing frameshifts (Figure 2C). The number of mutated genes unique to NSE tumors were 1707, while CGRP tumors harbored mutations in 107 unique genes. Also, 17 mutated genes were common between the two models (Figure 2D). It is known that not all SNPs are associated with cancer progression and that their impact may vary depending on the location of the SNPs within the genome. Accordingly, the high frequency of genetic variation including silent SNPs may not have a direct effect on tumor progression of NSE-p25OE but can still indicate crucial information about the genetic landscape of tumors. The direct impact of genetic variations identified in NSE/CGRP tumors is not yet fully understood and requires further investigation.

**Figure 2.**
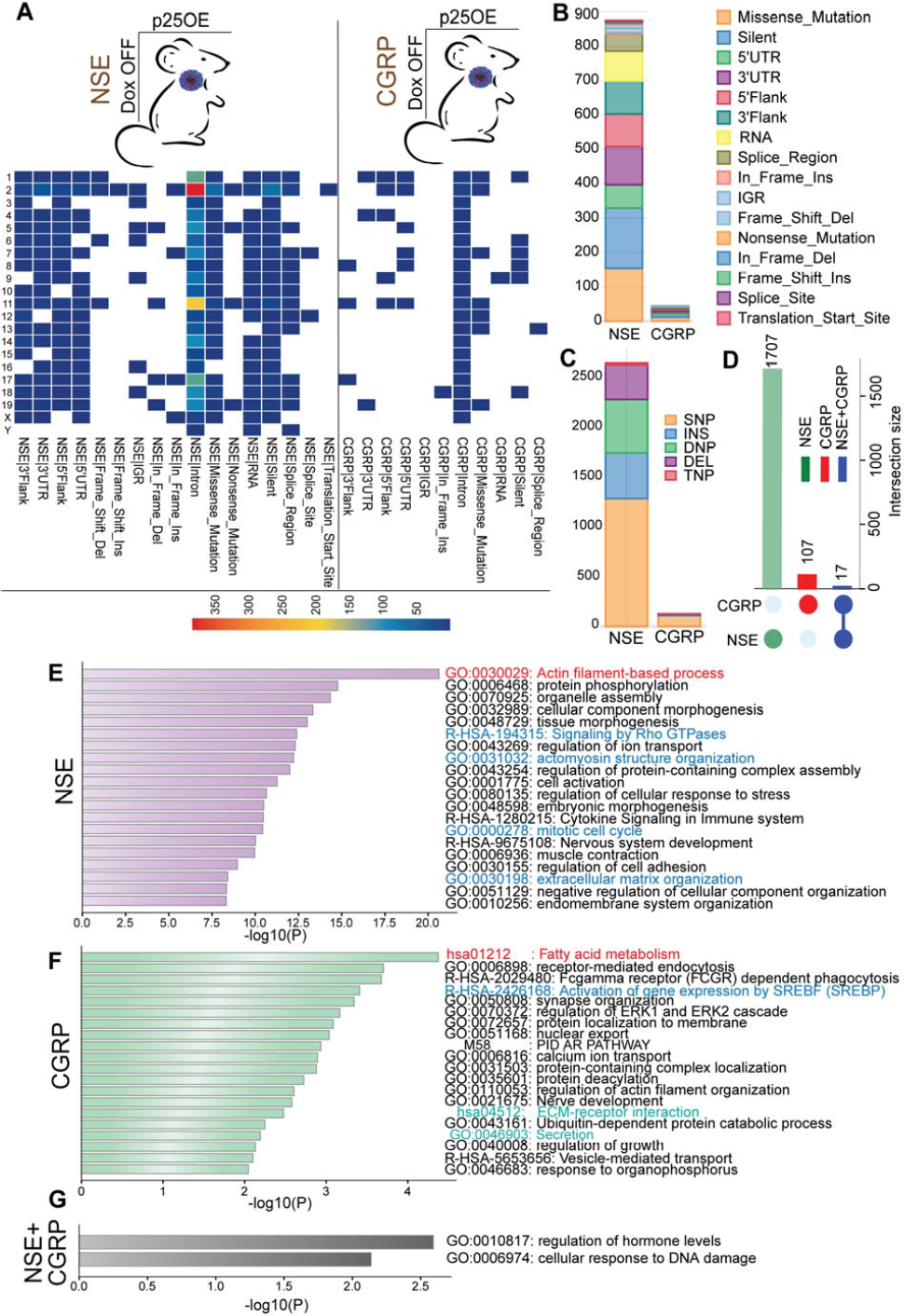
Mutational landscape of mouse tumor models. (A) Mutation profile of growing (Dox Off or p25OE) NSE (n=4) and CGRP (n=3) mouse tumors across chromosomes. (B) Plot of variant classification by type; frequency of variant (y-axis), colors denote types of variation. (C) Variant type presented as SNP (single nucleotide polymorphism), INS (insertion), DNP (double nucleotide polymorphism), DEL (deletion), TNP (triple nucleotide polymorphism); frequency (y-axis). (D) UpSet plot showing counts of unique or overlapping mutated genes in NSE and CGRP models. (E) Bar chart displaying enriched pathways associated with altered genes in NSE-p25OE mice; (F) Enriched pathways in CGRP-p25OE mice; (G) Enriched pathways common in –NSE and –CGRP p25OE mice. Significant functions are shown where p-value < 0.01.

To understand the molecular processes underlying tumorigenesis of NSE/CGRP models, pathway enrichment analyses of altered genes were performed. The most highly enriched mutated gene set in NSE-p25OE tumors was involved in ‘actin filament-based process’ (Figure 2E). Other altered cancer-related GO pathways included mitotic cell cycle, Rho-GTPases, cytokine signaling, cell adhesion, and extracellular matrix organization (Figure 2E). In addition, KEGG database indicated enrichment of phosphatidylinositol, GnRH (gonadotrophin receptor hormone), and oxytocin signaling including actin/cytoskeletal-based processes (Figure S1A). Dysregulation in the GnRH and oxytocin signaling pathways have been implicated in several cancers [27, 28], but unexplored in MTC, suggesting an important avenue for further investigation. In contrast, KEGG enrichment of mutated genes in CGRP-p25OE tumors displayed ‘Fatty acid metabolism’ as the most significantly altered pathway besides glutamate receptor clustering (Figure 2F and S1B). Of note, glutamate signaling is actively involved in bioenergetics and metabolic pathways in cancer [29]. In agreement, glutamate receptor antagonists are known to suppress MTC and carcinoid NET growth and metabolic activity [30]. Likewise, the activation of SREBP, a master transcription factor, and regulator of lipid metabolism [31] was altered in CGRP mice (Figure 2F). These data provide evidence of metabolic derangement in CGRP-p25OE mice. The overlapping mutated genes between NSE and CGRP were prominently enriched in the ‘regulation of hormone’, a characteristic of MTC patients showing high biosynthetic and hormone secretory activity [32]. In addition, ‘cellular response to DNA damage’ processes were identified in both NSE/CGRP models (Figure 2G). Based on previous reports, activation of DNA damage response genes is common in MTC whereby modification of chromatin machinery favors a drug-resistant phenotype [33, 34].

Heretofore, we showed that activation of aberrant Cdk5 develops human-like MTC that accumulates mutations altering cell cycle and metabolic profiles in respective models. However, it is critical to understand if the mutational landscapes of mouse tumors emulate their human counterparts. To evaluate the application of these tumor models as prototypes for human disease, we compared mutated genes in mouse and human tumors. Out of 1693 altered genes, 42 orthologs were common between NSE-p25OE and human MTC [35] (Figure 3A, S2A, and Table S2). The prominent genes intersecting between human and mouse MTC, such as RB1, AKT1, and NDGR2, are integral to processes including cell proliferation, cell cycle, and immune system function [36–38]. Several other common mutated genes including *SMARCA2, Cdk6, BTG3, ROCK1, PLD1*, and *DEFB1* play important roles in cancer progression but are unexplored in MTC and represent interesting targets for future investigation (Figure 3A). The top significantly enriched biological processes common between NSE-p25OE and human tumors include ‘G1/S transition of mitotic cycle’, and ‘mitotic cell cycle phase transition’ (Figure 3B and S2B-C). Moreover, pathways linked to mutated genes were highly clustered across FOXO3A signaling, NOTCH-NFkB signaling, and G1/S phase transition, consistent with those previously reported in MTC patients (Figure 3B) [39–41]. Conversely, CGRP-p25OE mouse vs. human tumor comparison revealed five intersecting genes harboring mutations in *RPS6KA2, GPM6A, PATZ1, HACD4*, and *CBFA2T3* (Figure 3C, S3A, and Table S2). Biological processes connected to altered genes were mainly involved in cellular metabolism (*i.e*., fatty acid metabolism, glycolytic process, and nucleotide metabolism). Notably, a significant impact on the biosynthesis of unsaturated fatty acids and fatty acid elongation pathways was recognized (Figure 3D and S3B-C). In concordance, impairments in fatty acid metabolism, purine metabolism, and tri-carboxylic acid cycle were reported in MTC patients compared to healthy controls [42]. In summary, our results demonstrated a profile of signaling pathways altered in human tumors and replicated in our mouse models. Of particular interest are the ‘mitotic cell cycle process’ in NSE-p25OE and ‘metabolic impairment’ in CGRP-p25OE models. Further in-depth understanding of these processes can uncover the underlying cause of the differential rate of tumor progression in individual models and possibly human patients.

**Figure 3.**
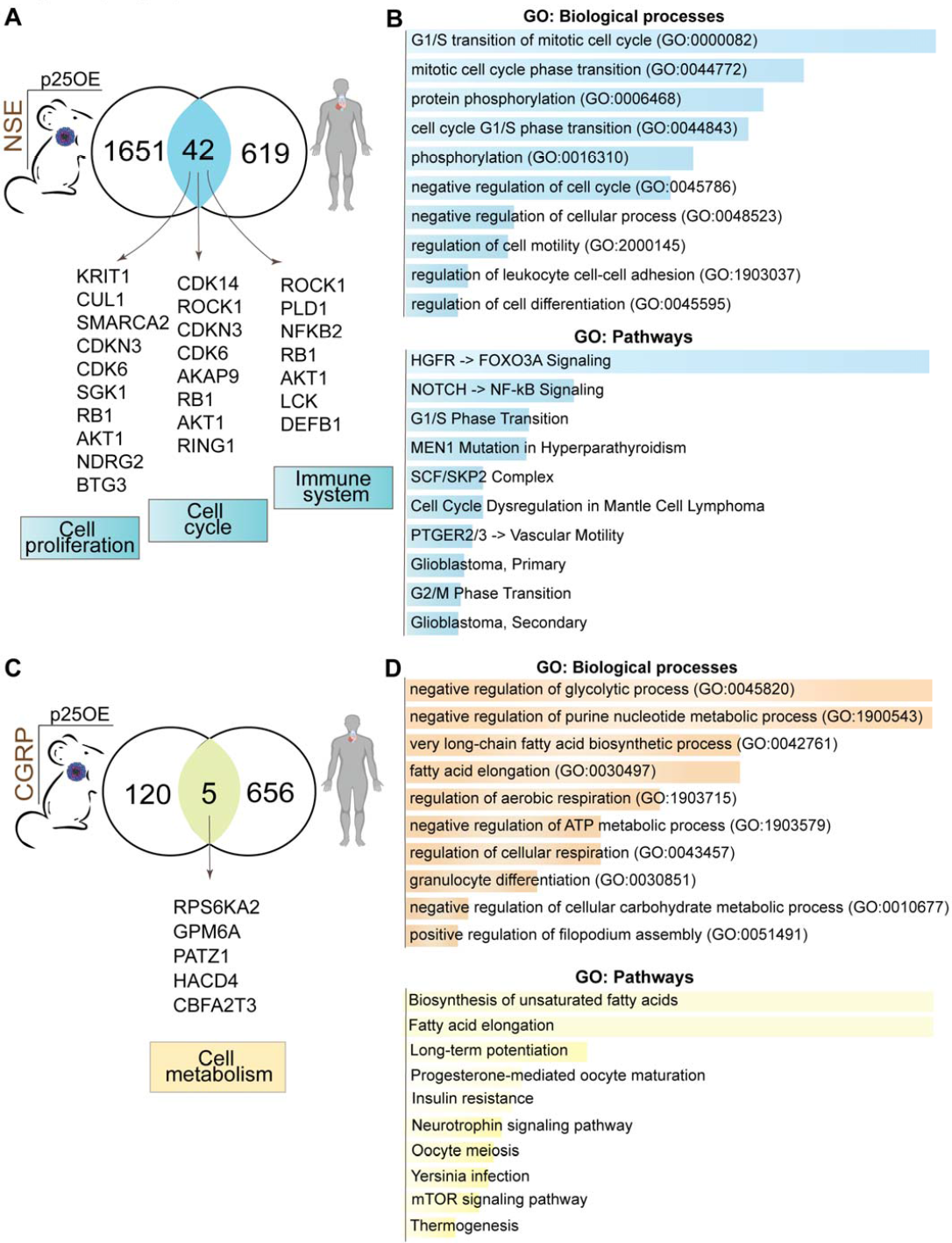
Comparison of mouse and human tumors mutational profile. (A) Venn diagram showing overlapping genes between NSE-p25OE mice and human tumors [35]. (B) GO term enrichment analysis shows biological processes and pathways associated with the altered genes shared by NSE-p25OE mice and human tumors. (C) Overlapping mutated genes in CGRP-p25OE mice and human tumors [35] are shown. (D) GO analysis of overlapping genes shared by CGRP-p25OE mice and human tumors. Bars sorted by p-value ranking; p-value cut-off < 0.05; GO, Gene Ontology.

### Functional impact of p25OE on transcriptomics of mouse tumors

Gene mutations have the propensity to induce transcriptional changes which influence cancer cell progression and responses to chemotherapy. To identify the alterations in gene expression, bulk RNA sequencing was performed on growing and arrested MTC tumors derived from NSE and CGRP models. The principal component analysis demonstrated distinct segregation of gene expression between growing and arrested tumors derived from NSE/CGRP (Figure 4A). Further gene expression analysis indicates a total of 4920 differentially expressed genes (DEGs), of which 2426 were upregulated and 2429 were downregulated in growing NSE-p25OE tumors. Whereas CGRP-p25OE revealed a total of 4348 DEGs, of which 2079 were upregulated and 2269 were down-regulated (Figure 4B). Intersection size of unique and common DEGs up- and down-regulated in NSE and CGRP tumors was determined by upset plots. Notably, both NSE/CGRP tumors showed upregulated transcripts of Cdk5R1 and Cdk5RAP2 indicating augmented transcriptional regulation of Cdk5 signaling components in these models (Figure 4C and Table S3).

**Figure 4.**
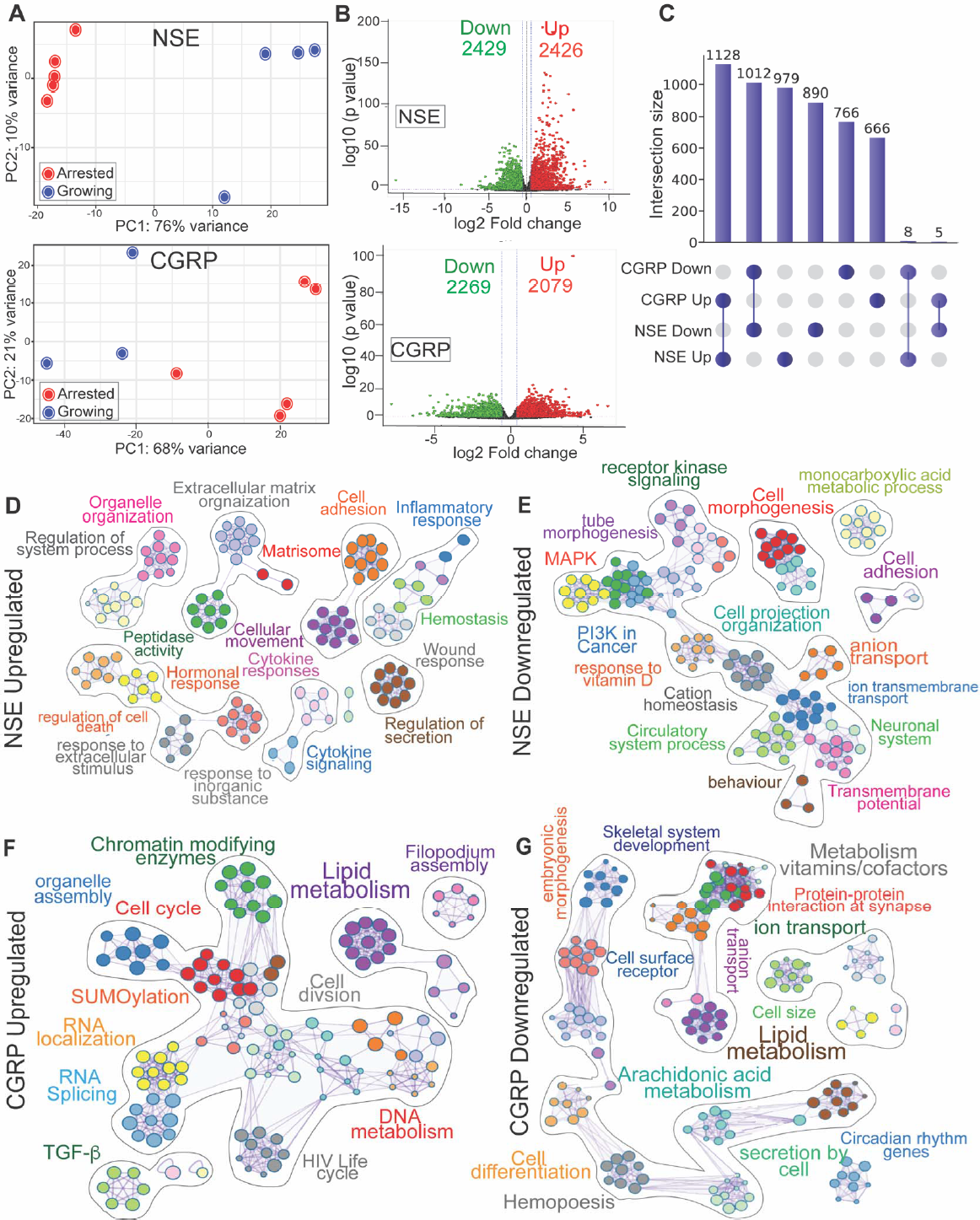
Transcriptomic analysis of mouse models. (A) Principal component analysis of RNA-seq reads in NSE-p25OE (n = 4 growing; n = 5 arrested) and CGRP-p25OE mice (n = 5 growing; n = 5 arrested). (B) Volcano plot presenting differentially expressed genes (DEGs) in growing vs. arrested tumors of NSE-p25OE and CGRP-p25OE models. Red = upregulated DEGs; Green = downregulated DEGs; log_2_FC>1.5. (C) UpSet plot summarize unique and overlapping DEGs in NSE and CGRP p25OE models. Visualization of enrichment network of DEGs up and downregulated (D-E) in NSE-p25OE mice, and (F-G) in CGRP-p25OE mice. Cluster annotation is indicated by color code, nodes with the same color are closely spaced and associated with the same cluster, clusters were labeled manually; enriched terms with a similarity score of >0.3 are connected by edges.

Pathway and process analyses performed on DEGs were clustered based on similarities of enriched terms (p-value < 0.01, gene count, and enrichment factor > 1.5). Network analysis of enriched nodes with a similarity score of >0.3 was connected by edges. Accordingly, upregulated DEGs in NSE tumors show hallmark clusters involved in processes such as cell adhesion, positive regulation of cell migration, cell motility, actin filament polymerization, cytokine signaling, T-cell activation, and lymphocyte proliferation (Figure 4D). The unique genes integrated within the main clusters are shown in Table S4. The downregulated DEG in NSE tumors clustered cell morphogenesis, ion transport, receptor kinases, MAPK, and PI3K activities (Figure 4E). Notably, processes such as tissue morphogenesis, ion transport, cytokine signaling, and extracellular matrix organization were afflicted both with mutations and transcriptional dysregulation in NSE tumors. Further, TRRUST database uncovered the transcriptional factors (TF) namely, c-Jun, Sp1, and NFkB as putative regulators of DEGs in NSE-p25OE (Figure S4A). Interestingly, the activity of c-Jun is regulated via Rho GTPases which in turn drives the transcription of genes involved in the cell cycle progression [43]. This aligns with the exome data showing altered Rho GTPase signaling in NSE mice (Figure 2E). Likewise, the Sp1 is a known regulator of ion transport in the thyroid cancer [44] while the enrichment of NFkB signifies a plausible contribution to cytokine and inflammatory responses in NSE-p25 tumors [45].

Analysis of upregulated DEGs in CGRP-p25OE revealed enrichment clusters across the cell cycle, cell division, metabolism of lipids, nucleotide metabolic processes, SUMOylation, RNA splicing/localization, chromatin-modifying enzyme, and transforming growth factor beta (TGFβ) (Figure 4F). The prominent gene hits involved in these processes are enlisted in Table S5. The downregulated transcripts altered many metabolic pathways like those afflicted by mutations in CGRP tumors (Figure 4G and Figure 2F). TRRUST query identified Ncoa1 as the putative transcriptional regulator of DEGs in CGRP tumors, which is shown to regulate lipogenic and glucose metabolic pathways [46, 47] (Figure S4B). These results indicate dysregulation of distinct pathways impacted both by mutations and gene expression changes downstream of hyperactive Cdk5 in NSE/CGRP models of MTC.

### Consequence of somatic mutations on gene expression of mouse tumors

Genetic abnormalities can influence gene expression and signaling pathways contributing to the tumorigenic process. To prioritize the genes that confer a growth advantage in our MTC models, we determined the influence of mutations on the expression level of their residing genes. The upset plot shows 128 and 144 altered genes that correlate with changes in mRNA expression of NSE-p25OE tumors (Figure 5A and Table S6). The top-upregulated mRNA gene sets hit by somatic mutations were enriched in spindle assembly, APC/C-CDC20 complex, kinetochore assembly, and NOTCH signaling pathways (Figure 5B). Conversely, the key mutated genes that correlated with downregulated transcripts were primarily involved in cancer-associated proliferative signaling (Figure 5C). Subsequent prioritization of mutated genes based on changes in gene expression displayed BUB1B, MAD1L1, and DDL4 as main targets, elevated in growing tumors compared to arrested tumors signifying a potential tumor-promoting role (Figure 5D). In addition, the suppression of EPHA4 and NFkB2 mRNA levels in growing tumors indicates tumor suppressive function of the residing mutations (Figure 5E).

**Figure 5.**
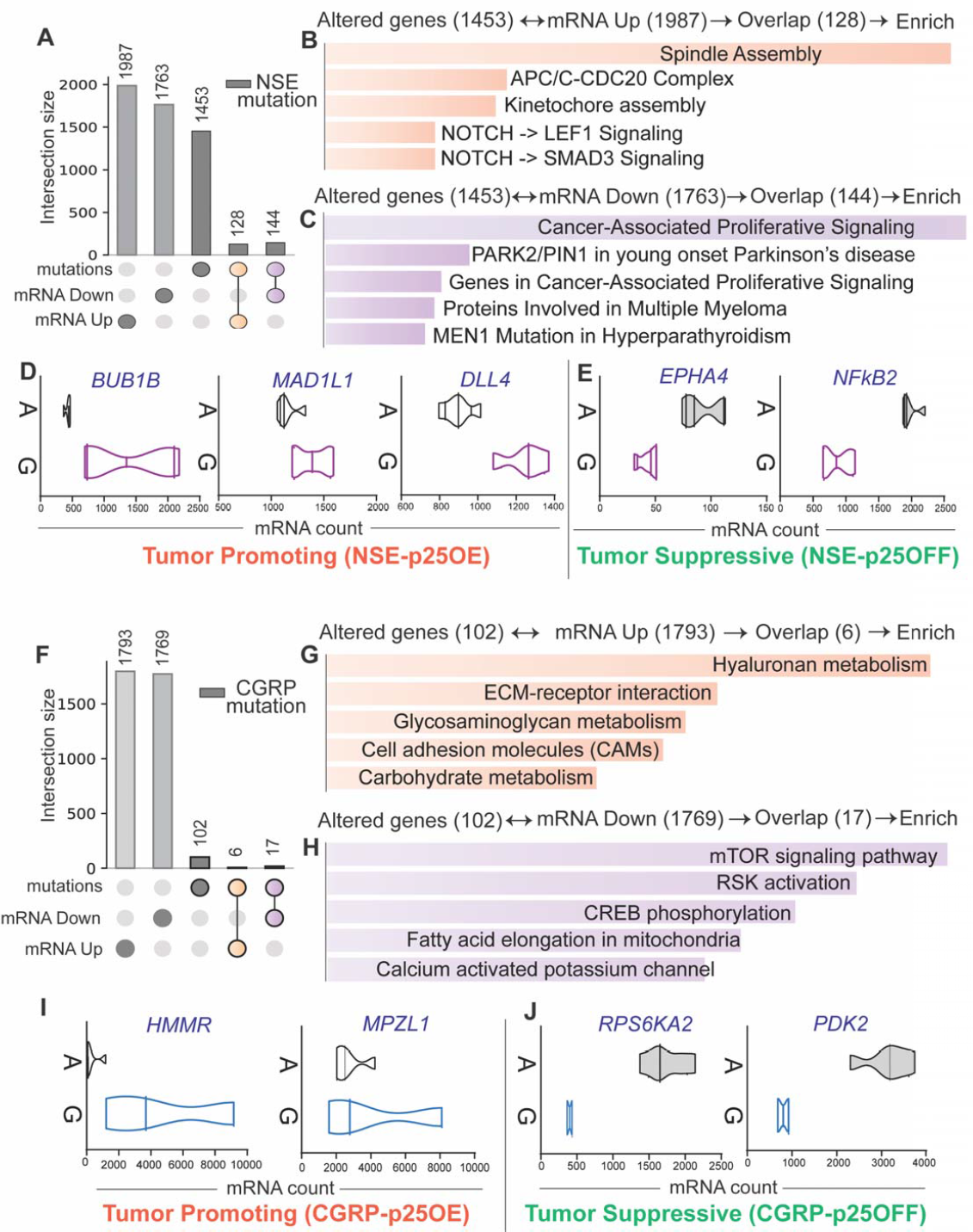
Impact of genetic alterations on mRNA expression. (A) UpSet plot depicts the overlap between mutations and corresponding gene expression in NSE-p25OE mice. (B-C) Pathway visualization of overlapping genes afflicted both with mutations and changes in mRNA expression. Bar charts showing pathways associated with the mutated genes where mRNA was up-(B) or down-regulated (C) in NSE-p25OE mice. (D) Violin plots compare mRNA reads of mutated genes involved in spindle assembly and Notch signaling. (E) Violin plots compare mRNA reads of mutated genes enriched in cancer-associated proliferative signaling (G, growing; A, arrested tumors). (F) Intersection size of mutated genes and corresponding mRNAs in CGRP-p25OE mice. (G-H) Functional enrichment of overlapping mutated genes where mRNA was either up-(G) or down-regulated (H) in the CGRP-p25OE mice. (I) Plots showing mRNA reads of mutated genes associated with metabolic pathways and cell migration, or (J) mRNA reads of mutated genes involved in mTOR signaling.

Next, we determined the consequence of mutations on the expression profile of the CGRP-p25OE model. Overlap of WES with RNA seq data revealed six alterations that caused mRNA upregulation (Figure 5F and Table S6). Pathways analysis disclosed metabolic processes such as hyaluronan metabolism, glycosaminoglycan metabolism, and carbohydrate metabolism in addition to extracellular matrix-receptor interaction and cell adhesion processes associated with upregulated mutated genes (Figure 5G). Further, 17 mutated genes correlated with transcriptional repression (Figure 5F). Enrichment analysis of intersecting genes indicates suppression of mTOR signaling, RSK activation, CREB phosphorylation, and fatty acid elongation in mitochondria (Figure 5H). Finally, gene prioritization based on alterations in gene and transcript levels revealed HMMR, and MPZL1 as tumor promoters, whereas RPS6KA2 and PDK2 as tumor suppressors in growing CGRP tumors (Figure 5I-J). In conclusion, the comparative mutational and transcriptomic landscape of CGRP/NSE models revealed candidate genes that may serve as potential regulators of tumor progression downstream of aberrant Cdk5.

### Histodiagnostic characterization of Cdk5 activity in mouse and human tumors

We previously showed that hyperactive Cdk5 plays a critical role in MTC progression [10]. A positive correlation was demonstrated between Cdk5 and its downstream targets in human MTC. The main protein phosphorylation substrates of this aberrantly active kinase included P-T143 FAM53C, P-T202 LARP6, P-S988 RBL1, P-S17 H1.5, and P-S391 SUV39H1 [48]. Both NSE/CGRP models manifest aberrant Cdk5 activation as indicated by increased Cdk5-dependent phosphorylation in growing (p25OE) versus arrested tumors (p25OFF) (Figure. S5A), suggesting these phosphosites may serve as biomarkers for the detection of Cdk5-driven human tumors. However, the polyclonal antibodies first used to detect these Cdk5 targets were raised against short phospho-peptide epitopes that were limited in specificity and often detected cross-reactive proteins harboring similar phosphorylation site motifs. Considering the potential importance of aberrant Cdk5 in MTC diagnosis, we sought to derive more selective and specific monoclonal antibodies (mcAb) that could precisely probe Cdk5 activity in mouse and human tumors. Recombinant phosphorylation state-specific mcAb were generated in collaboration with the National Cancer Institute’s Antibody Program via its Antibody Characterization Laboratory (ACL). The specificity of mcAb in growing (G) and arrested (A) tumors were evaluated for their detection of Cdk5 phosphorylation sites by immunoblotting, immuno-cyto, and -histochemistry. Expression analysis of P-RBL1, P-LARP6, P-H1.5, P-SUV39H1, and P-FAM53C showed improved efficiency and reduced non-specific binding of mcAb over polyclonal (pcAb) (Figure S6A-E). Further, the mcAb was tested on mouse cells derived from NSE-p25OE cells. These cells preserve the characteristics of mouse MTC where cell proliferation is regulated by p25OE (Figure 6A). Increased nuclear staining of Cdk5 target sites– P-RBL1, P-LARP6, and P-H1.5 were identified in growing/p25OE mouse cells compared to their arrested/p25OFF counterparts (Figure 6B). A similar effect was mirrored in CGRP/NSE-derived mouse tumor tissues where the expression of phospho-targets was higher in p25OE vs p25OFF tumors, suggesting dependency of these phosphosites on aberrant Cdk5 activity (Figure 6C). The improved detection of aberrant Cdk5 activity in inducible/arrestable mice models spurs the efforts to evaluate these mcAbs in human cells and tissues. Human MTC TT cells known for Cdk5-dependent cell growth were utilized was probing key phosphoproteins [49]. The main phosphoproteins including, RBL1 (RB transcriptional corepressor like 1) regulate the cell cycle, and proliferation by modulating chromatin structure. LARP6 (La Ribonucleoprotein 6), a translational regulator shuttles between cytoplasm and nucleus promotes nucleic acid binding while linker histone H1.5 facilitates chromatin compaction. The immunocytochemical staining of TT cells displayed nuclear localization of these phosphoproteins similar to that observed in mouse MTC cells, consistent with the known function of these proteins in chromatin structure modulation. Having established selective detection of these aberrant Cdk5 effectors in human cells, we obtained histological tumor sections from a small cohort of MTC patients to assess the presence of these sites. Interestingly, the immunohistochemical results showed varying degrees of phospho-site staining across MTC patients confirming a characteristic heterogeneous expression profile of MTC tumors (Figure 6E).

**Figure 6.**
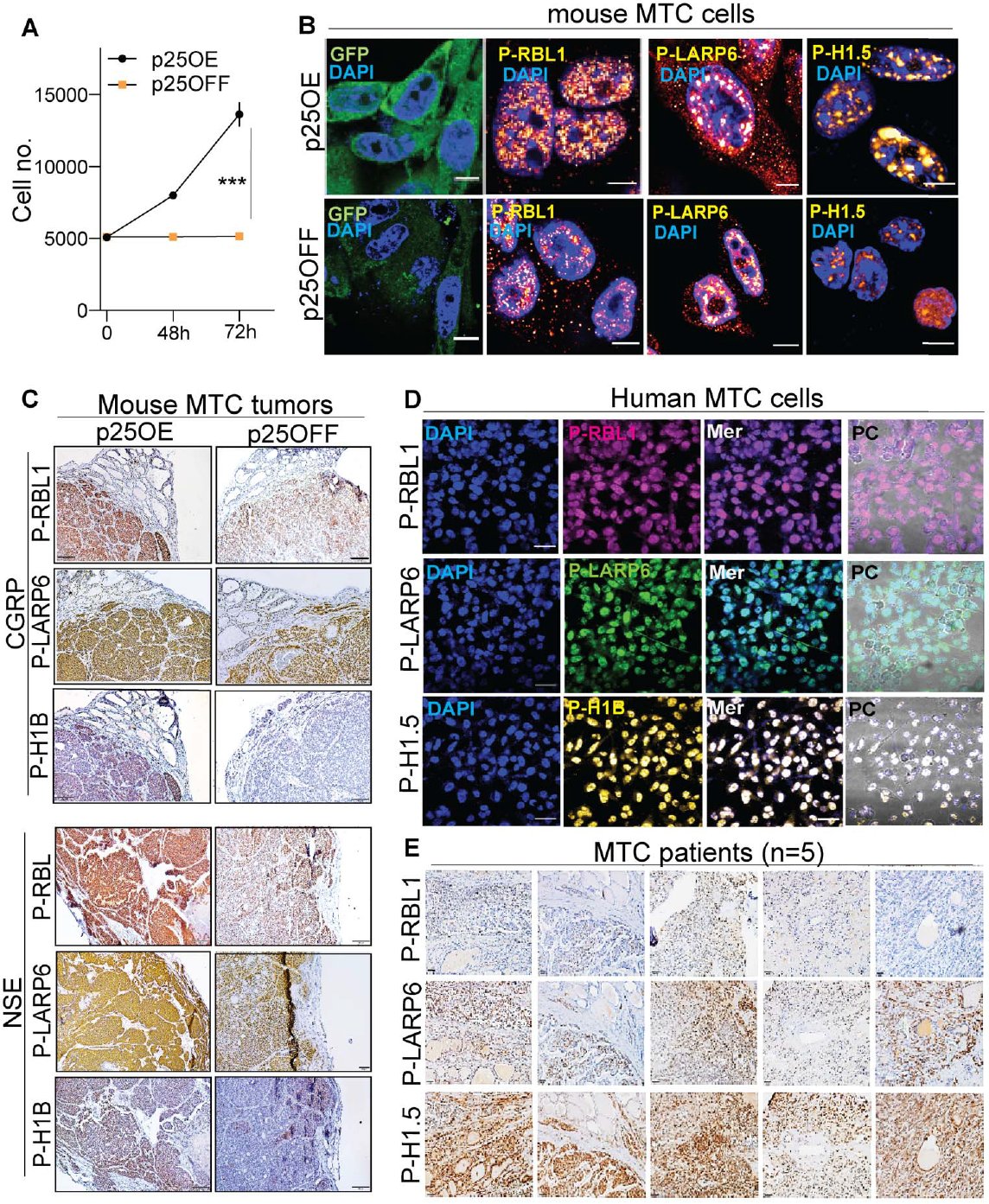
Efficacy of monoclonal antibodies for detection of aberrant Cdk5 activity. (A) Graph showing cell proliferation in mouse MTC cells under p25OE vs. p25OFF conditions, n = 3, Mean ± SD, *** p < 0.001. (B) Immunocytochemical analysis of Cdk5-dependent phosphorylations in mouse MTC cells as indicated, scale; 70 μm. (C) Immunohistochemical staining of Cdk5 phosphorylations in growing (p25OE) and arrested (p25OFF) tumor sections derived from NSE/CGRP models, scale; 50 μm. (D) Immunofluorescent and (E) Immunohistochemical micrographs of Cdk5 phosphosites in human TT cells and patient tumor sections, respectively (n = 5), scale; 50 μm.

Our data suggest that these phosphorylation biomarker detection reagents have the potential to identify aberrant Cdk5-driven human patient tumors. To explore this further, the expression of key Cdk5 phosphorylation sites including P-RBL1 and P-LARP6 was more broadly assessed across the three independent human MTC tissue microarrays (TMA 1-3), and compared with the normal non-tumor controls such as the prostate, placenta, spleen, and liver (Figure 7A-B). Quantification of phospho-site expression indicated by optical density (OD), shows 27% and 44% of patients with elevated P-RBL1 and P-LARP6 in TMA1 respectively while 28% of patients exhibited an increase in both P-LARP6/P-RBL1 (n=25; TMA1; stages: pT1bNX, and pT2N0). In TMA 2, 25% of patients with lymph node metastasis showed increased P-RBL1/ P-LARP6 (n=12; stage: pT3N1b). Quantification of TMA 3 displayed an increase of both phosphosites in ∼36% of patients (n=11; stage: pT2NX and few cases with metastatic deposits in lymph nodes) (Figure 7C-D). Overall histo-analysis of tissue microarrays showed a significant increase of Cdk5 phosphosites in patient tumors trending towards metastasis compared to the normal tissues (Figure 7A-D). Until last year, the World Health Organization (WHO) did not recommend the testing of Ki-67 proliferation index in cases of MTC. Therefore, it is advised by the pathologist in this study to acknowledge the fact that our dataset cases were not tested for Ki-67 as it was not considered a standard practice during the time of surgery and pathological diagnosis. In summary, our findings characterize the efficacy of mcAbs in identifying aberrant Cdk5 activity across mouse and human tumors manifesting diagnostic value in stratifying cancer patients that are most likely to have clinical benefits from Cdk5 inhibitors.

**Figure 7.**
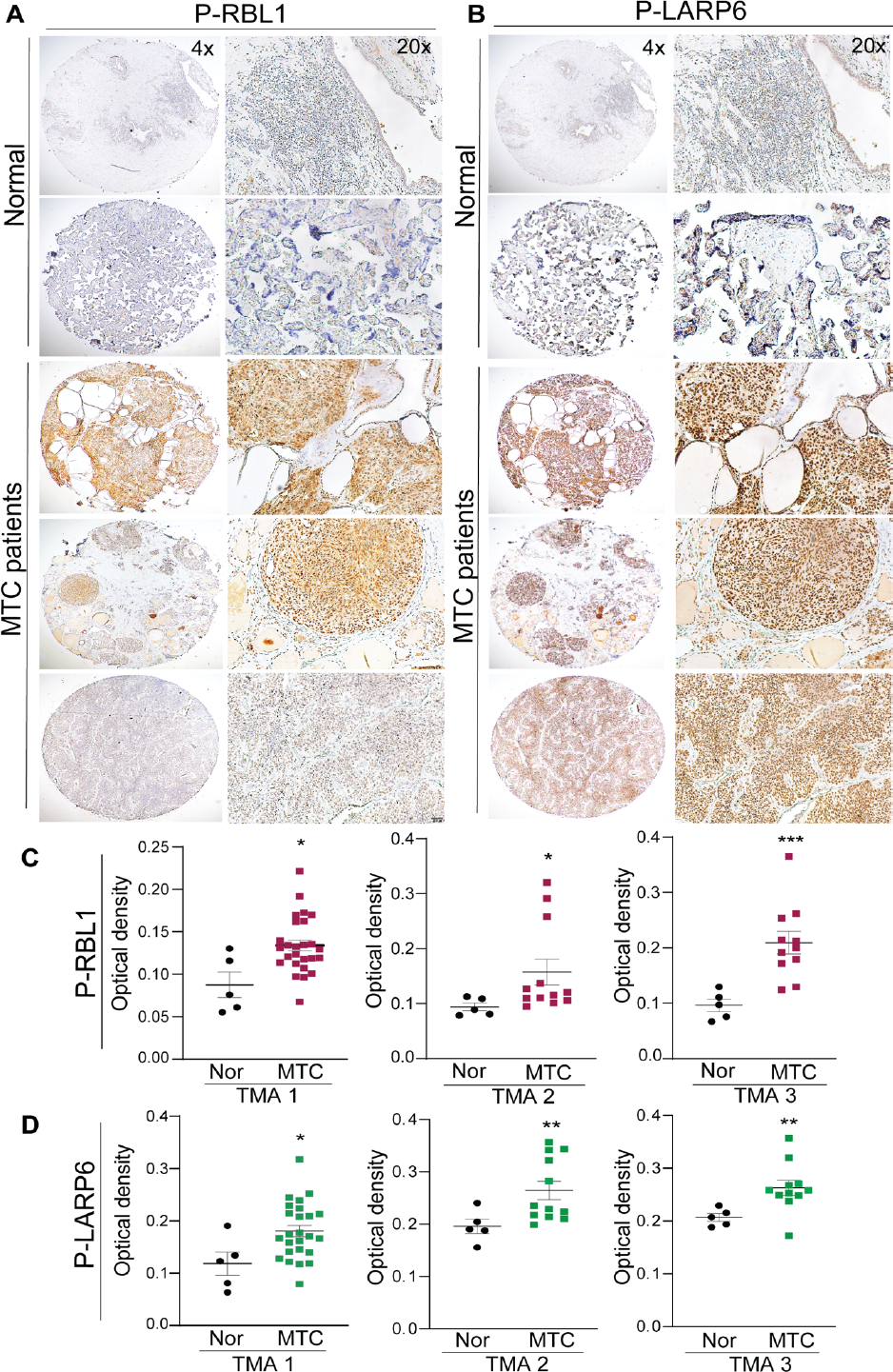
Histological evaluation of aberrant Cdk5 activity in human tissue microarray. (A-B) Representative immunohistochemical micrographs showing Cdk5 phosphosites *viz.* P-RBL1 and P-LARP6 in tissue microarray sections of MTC tumors. magnification; 4x (scale, 200μm), 20x (scale, 50 μm). (C-D) Quantification is presented as the optical density for (C) P-RBL1 (TMA 1-3), and (D) P-LARP6 (TMA 1-3). TMA 1: Nor = 5, tumor = 25; TMA 2: Nor = 5, tumor = 12; TMA 3: n = 5, tumor = 11. *p < 0.05, **p < 0.01, ***p < 0.001; values compared using Student’s *t-*test with Welch’s correction.

## Discussion

The lack of reliable animal models that mimics aggressive MTC disease is a major setback in the investigation of genetic alterations, identification of diagnostic biomarkers, and molecular mechanisms associated with malignant growth. Cdk5 is aberrantly activated in several NETs including MTC, contributing to tumor development. To circumvent the lack of appropriate models, we utilized Cdk5 as a tool to generate conditional transgenic mice that develop ‘slow’ and ‘rapid’ onset human-like MTC tumors. Aberrant Cdk5 can be characterized by the simultaneous activation of multiple pathways regulating cell proliferation and invasion. Hence it is critical to capture the complexity of signal transduction downstream of hyperactive Cdk5 to identify distinctive markers of aggressiveness and targets for therapeutic intervention. Here we characterized two mouse models of MTC, namely NSE-p25OE and CGRP-p25OE, mediating mild and aggressive onset of the disease. To understand the genes and pathway perturbation in these models, exomic and RNA sequencing was performed in tumors derived from the respective models. Computational analysis identified potential pathways associated with the altered genes in mouse tumors and those intersecting with human tumors. In addition, critical genes were prioritized based on changes in the gene and transcript levels.

Major pathways associated with altered genes in NSE-p25OE tumors were involved in actin filament-based processes. Actin-dependent enrichment included mitotic cell cycle, cell adhesion, Rho GTPase, actomyosin, and extracellular matrix organization, processes known for their role in cancer cell proliferation [50–53]. In agreement, the causal impact of unbalanced actin dynamics in MTC tumor invasiveness and growth has been suggested [54]. Cdk5 is known to regulate actin microtubule cytoskeleton, suggesting a Cdk5-dependent phenotype is acquired by the NSE model [55, 56]. Furthermore, the main overlapping mutated genes and pathways altered in NSE-p25OE and human tumors were comprised of RB1, AKT1, SMARCA2, PLD1, FOXO3a, Notch, NFkB, and G1/S phase transition. Notably, many of these pathways have previously demonstrated oncogenic or tumor-suppressive roles in thyroid cancer [36, 39, 40, 57]. Moreover, network pathway analysis of transcriptomic data revealed recurrent pathways afflicted both by mutations and gene expression changes. These data suggest that neoplastic transformation in the slow growing NSE-p25OE model recapitulates the indolent form of human disease that may have a higher plausibility of dysregulated actin dynamics and mitotic cell cycle processes.

Conversely, the rapidly growing CGRP-p25OE model harbored mutations that predominantly impacted fatty acid metabolism. This finding is supported by a recent study that demonstrated perturbation of fatty acid metabolism in MTC patients compared to healthy controls [42]. In addition, the overlapping mutated genes between CGRP-p25OE mouse and human tumors clustered pathways associated with fatty acid biosynthesis. This highlights the likelihood of metabolic dysregulations in instilling aggressive phenotype in CGRP mouse and human MTCs. An increasing number of studies show that malignant tumors are highly dependent on lipid metabolism and fatty acid synthesis for growth and survival [58, 59]. Cdk5 is an emerging candidate entangled in several metabolic conditions including cancer, diabetes, and obesity [60] [61]. Notably, Cdk5-dependent phosphorylation of PPARγ and PRKAG2 impairs key metabolic sensors such as adipsin, adiponectin, and AMPK kinase [12, 62]. A direct connection between Cdk5 and lipid metabolism was recently reported where Cdk5-mediated phosphorylation of acetyl-CoA synthetase 2 (ACSS2) regulates lipid production and promotes glioblastoma growth [63].

Following WES, transcriptomic analysis of CGRP-p25OE tumors also revealed alteration in metabolic pathways including lipid metabolism, arachidonic acid metabolism, DNA metabolism, and metabolism of vitamins/ cofactors. Apart from metabolism, cell cycle, and cell division mRNA clusters were enriched in CGRP tumors. Periodic expression of cell cycle regulatory genes facilitates tumor cell proliferation [64]. For example, altered and periodic expression of cell cycle regulatory genes such as *PTTG1, AURKA*, or loss of *CDK/RB*, *p18*, and *p27* are known for promoting aggressiveness in MTC [65, 66]. Based on our analysis, we believe dysregulation of lipid metabolism and cell cycle processes in orchestration with aberrant Cdk5 may lead to aggressive phenotype in CGRP-p25OE tumors. Further investigation will be needed to validate these mechanisms.

We also examined the transcriptional correlates of mutated genes to prioritize crucial genes in mice models. The principal mutations that induced changes in the mRNA expression of NSE-p25OE were in the leading edge of mitotic spindle assembly and Notch signaling components. Our findings, in concordance with previous studies, suggest that abnormal mitotic spindle and chromosomal instability can be vital drivers in MTC progression [67] [68]. The involvement of Notch in the development of thyroid C-cells and MTC growth further corroborate our analysis [40]. The spindle assembly and Notch-related genes including *BUB1B, MAD1L1,* and *DLL4*, showed increased expression in growing tumors and, hence, may be viewed as tumor-promoters, whereas downregulated expression of mutated EPHA4 and NFKB2 in growing tumors as tumor suppressors.

Assessment of mutational impact on gene expression of CGRP-p25OE mouse revealed perturbation in metabolic and mTOR signaling pathways. Clearly, the disruption of metabolic pathways was consistent across the exome-sequencing and transcriptomic data, suggesting the need for functional investigation of metabolic targets in aggressive tumors. For instance, *HMMR* (hyaluronan-mediated motility receptor) is a putative neoplastic marker and a functional component of metabolic pathways associated with cancer progression and poor clinical outcomes, was found highly elevated in growing CGRP-p25 tumors [69]. Further, mutations and concomitantly decreased expression of RPS6KA2 (Ribosomal protein S6 kinase A2) and PDK2 (Pyruvate dehydrogenase kinase 2) implicate their function as potential tumor repressors in CGRP tumors. Of note, both RPS6KA2 and PDK2 are components of PI3K/Akt/mTOR pathway involved in cell cycle regulation and metabolic sensing in glycolytic cancers [70] [71] [72].

Alterations at the genetic and transcriptional levels of certain genes within a pathway can influence other genes eliciting interactions of multiple pathways. Our findings reinforce the need to classify patients based on aberrant Cdk5 activity, and further subclassify them into mild or aggressive forms based on the activation of distinct molecular pathways as described here. Here we developed and demonstrated new biomarker detection agents in the form of recombinant monoclonal antibodies that can distinguish Cdk5-dependent tumors. Previously, we showed that these biomarkers can predict anti-Cdk5 therapy responsiveness in patient-derived xenografts [48]. In the future, we aim to develop a monoclonal antibody-based diagnostic assay that can effectively stratify patients dependent on aberrant Cdk5 activity, as a potentially important step toward personalized therapy. Understanding the pattern of candidate oncogenic or tumor suppressor markers downstream of Cdk5-dependent MTC is crucial. Our findings meaningfully contribute to delineating the causes of malignant progression and can provide a path to potentially advance the effectiveness of the current treatment regimens.

In summary, we established transgenic mouse models that recapitulate both slow and rapid-onset human-like MTC tumors driven by aberrant Cdk5 activity. The CGRP-p25OE model mimics aggressive tumors primarily characterized by alterations in metabolic pathways. NSE-p25OE mice develop slow-growing tumors characterized by dysregulation of mitotic spindle assembly and Notch signaling components. Our findings encourage functional validation of putative targets that may lead to the development of personalized therapeutic strategies involving Cdk5 inhibitors in combination with patient-specific signaling modulators such as those targeting fatty acid metabolism or cell cycle regulators.

## Supporting information

Data summary of mutational profile of mouse tumors, related to Figure 2.

Summary of overlapping mutated genes in mouse and human tumors, related to Figure 3.

List of differentially expressed genes (DEGs) in NSE-p25OE and CGRP-p25OE mice, related to Figure 4.

List of gene hits associated with the main clusters of upregulated DEGs in NSE-p25OE and CGRP-p25OE tumors respectively, related to Figure 4.

List of gene hits associated with the main clusters of upregulated DEGs in NSE-p25OE and CGRP-p25OE tumors respectively, related to Figure 4.

List of altered genes that correlate with changes in mRNA expression of mouse tumors, related to Figure 5.

## Acknowledgments

We thank the National Cancer Institute Antibody Characterization Laboratory, the Office of Cancer Clinical Proteomics Research, and the Frederick National Laboratory for Cancer Research for the generation of recombinant monoclonal antibodies. We thank the UAB O’Neal Comprehensive Cancer Center Small Animal Imaging Facility, UAB Genomics Core, and the UAB High-Resolution Imaging Facility (P30CA013148-49, and S10OD028498). This research was supported by funding from the Robert Reed Foundation (H.C., S.R.), the International Thyroid Oncology Group (H.C.), the SDHB Pheo Para Coalition (J.A.B.), and the Neuroendocrine Tumor Research Foundation (J.A.B.). NIH awards that contributed to this work include the UAB Medical Scientist Training Program T32 GM008361 (B.H.), F31CA260945 (B.H.), K08CA234209 (B.R.), CA245580 (R.J.S.), American Cancer Society Postdoctoral Fellowship (A.M.C.), and ITOG (H.C.). Some aspects were facilitated by MH116896 and MH126948 (J.A.B.). Research was also supported by the UAB Center for Clinical and Translational Sciences Training Grant (TL1 TR001418).

## Author contributions

Conceptualization: P.G., J.A.B.; Methodology: P.G., J.A.B.; Investigation: P.G., B.H., N.K., R.T., F.W., P.K.; A.M.C.; D.L; Resources: H.C., B.R., R.J-S., S.R.; Writing Original Draft: P.G.; Review & Editing: J.A.B.; Funding Acquisition and Supervision: S.M., R. J.-S., S.R., J.A.B.

## Supplementary legends

**Figure S1.**
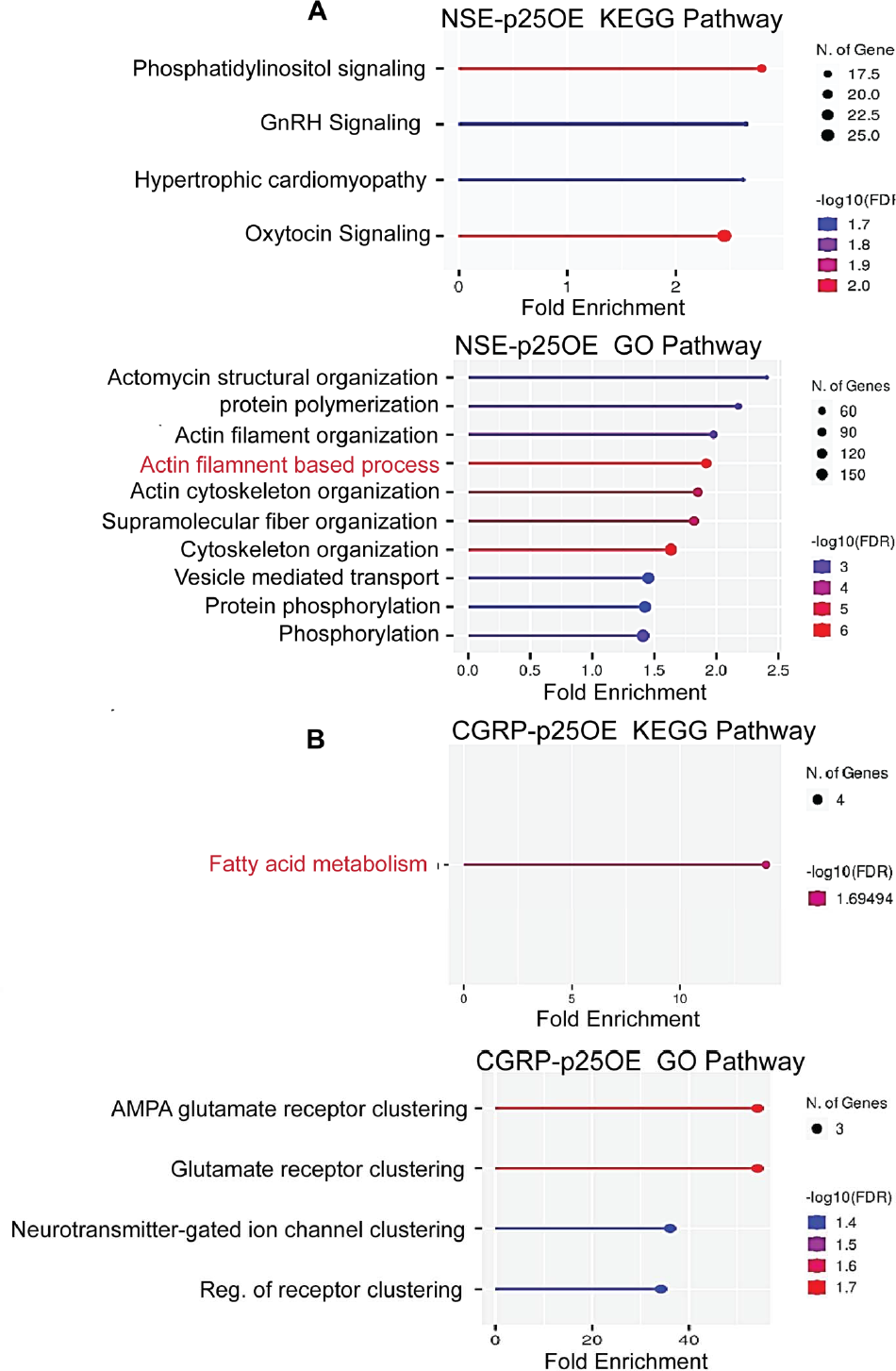
Enrichment analysis of mutated genes in mouse tumors. KEGG and GO enrichment in (A) NSE-p25OE and, (B) CGRP-p25OE mice. y-axis = pathway description; x-axis = fold enrichment. Analysis performed in Shiny GO 0.76.2; the bubble size indicates the number of genes, and the bar color code signifies the corrected p-value as indicated. GO: Gene Ontology (biological process); KEGG: Kyoto Encyclopedia of Genes and Genomes.

**Figure S2.**
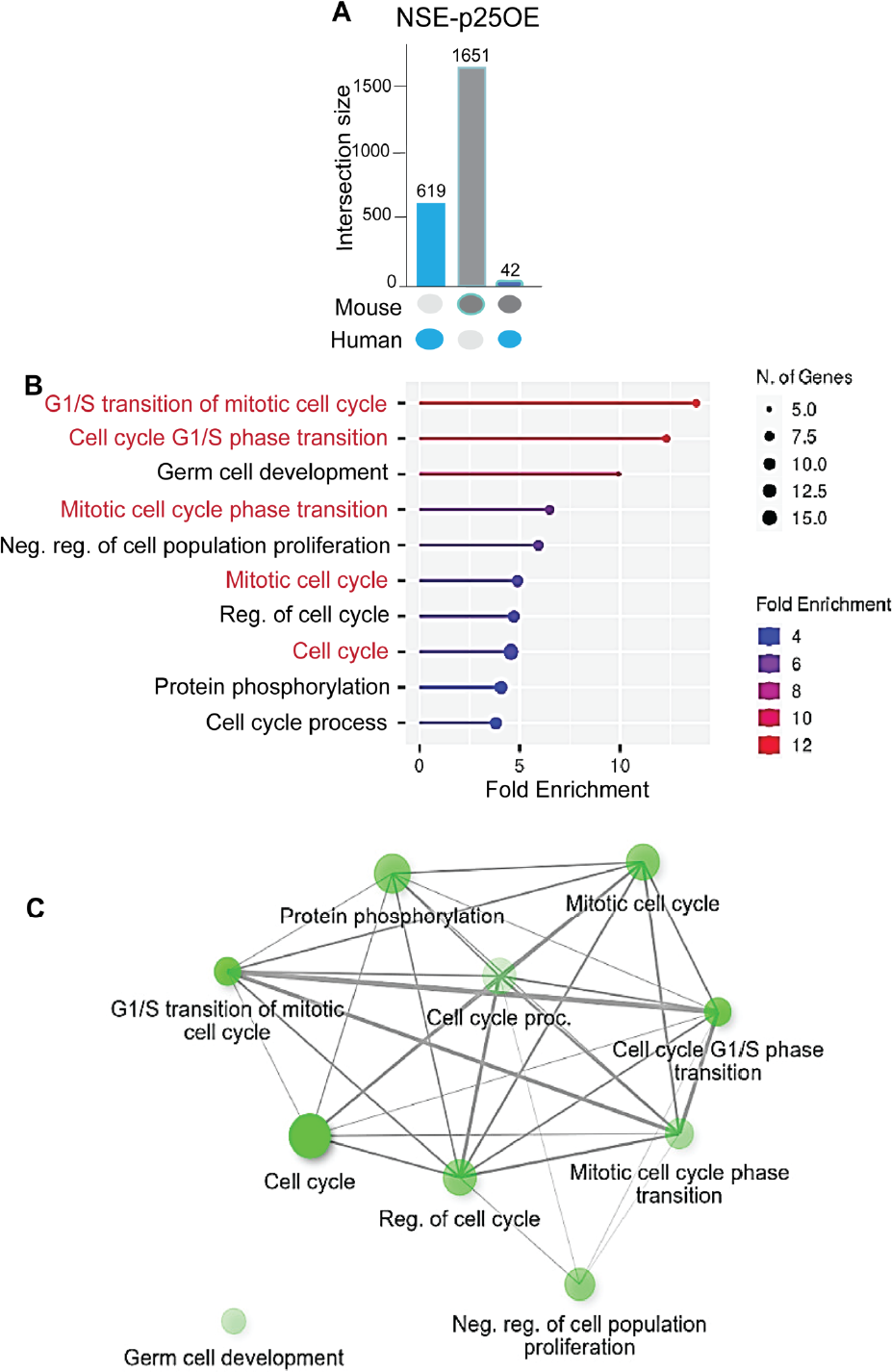
Mitotic cell cycle pathways shared by NSE mouse and human tumors. (A) UpSet plot showing intersecting mutated genes in mouse and human tumors. (B) Lollipop chart and (C) network visualization summarize significantly enriched GO terms common in NSE and human tumors. In a network diagram, two nodes are connected if they share >20% of their genes, FDR cutoff = 0.05.

**Figure S3.**
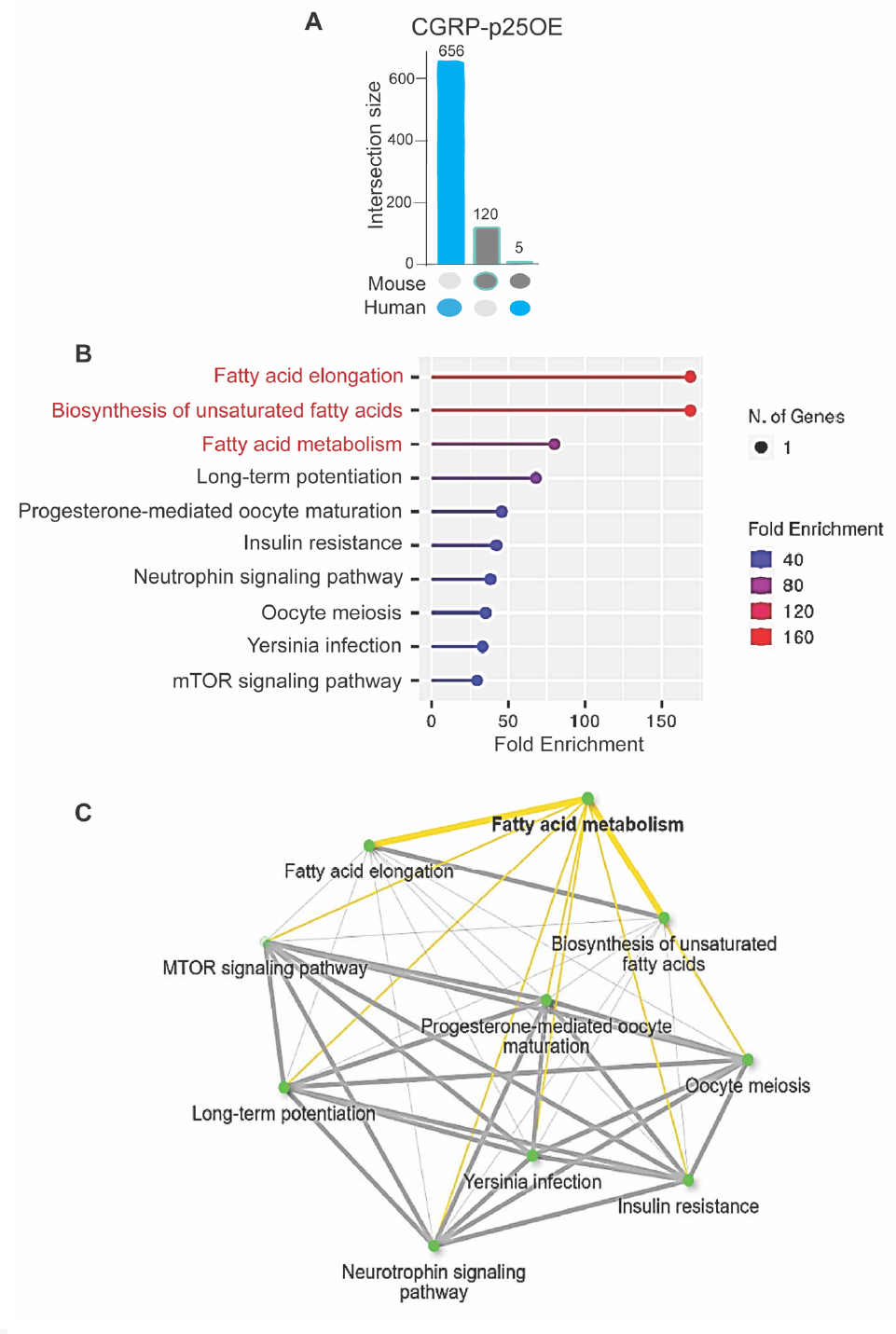
Metabolic pathways shared by CGRP mouse and human tumors. (A) Plot showing counts of unique and overlapping genes between mouse and human tumors. (B) Pathway and process enrichment and (C) Network tree of significantly enriched gene sets common in CGRP-p25OE mouse and human MTC, FDR cutoff = 0.05.

**Figure S4.**
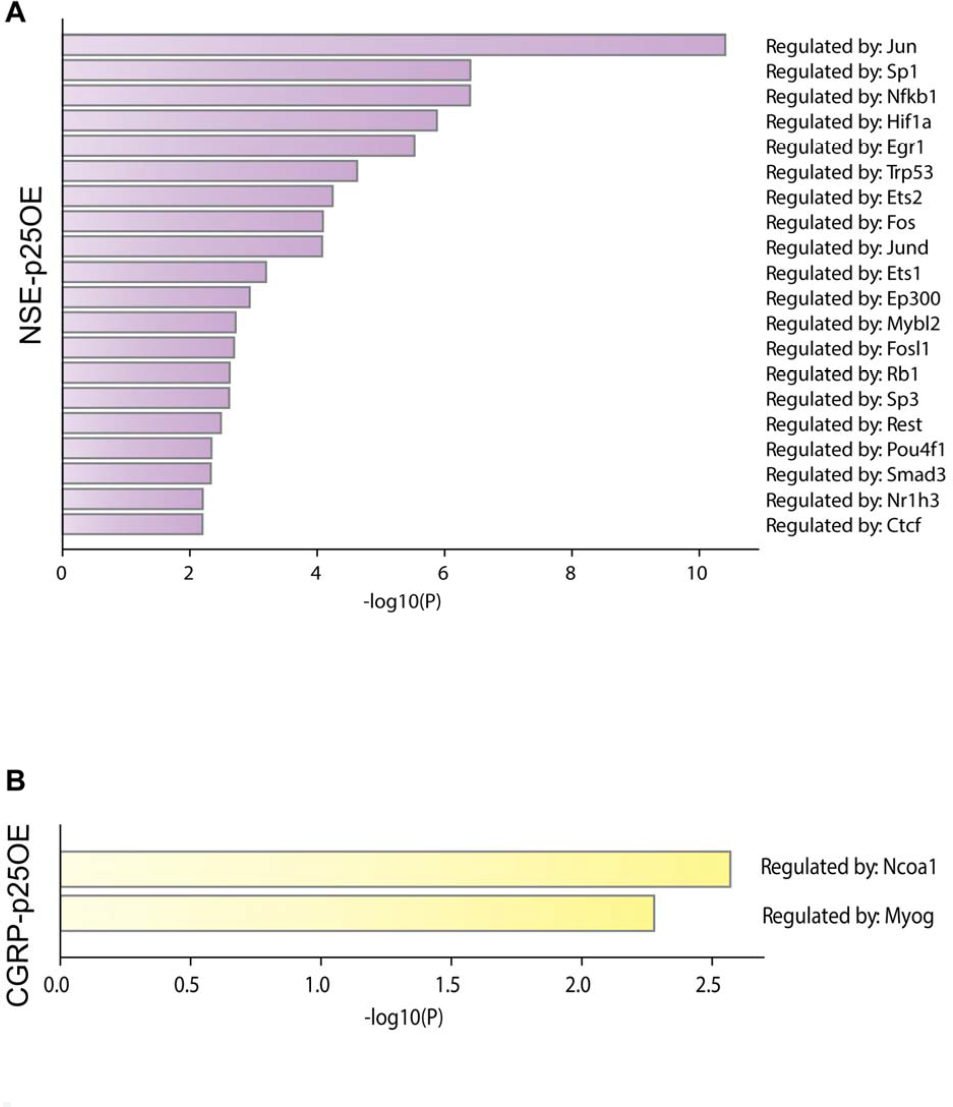
Enrichment of transcription factors in NSE-p25OE mice. (A) Plot showing putative regulators of DEGs in NSE-p25OE mice determined via TRRUST database [73]. (B) Summary of the enriched transcription factor in CGRP-p25OE mice. TRRUST database revealed putative regulators of DEGs in CGRP-p25OE mice. Terms with a p-value < 0.01; minimum count of 3, enrichment factor > 1.5 [74].

**Figure S5.**
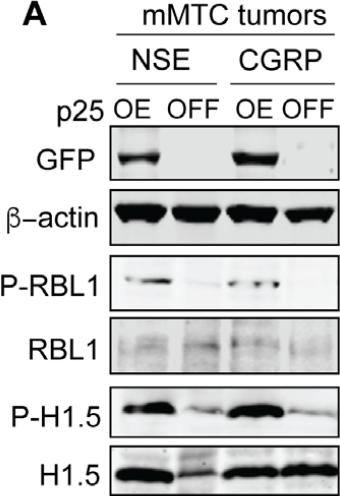
Detection of Cdk5 activity in NSE and CGRP tumors. (A) Immunoblots showing expression levels of p25GFP, P-RBL1, and P-H1.5 in tissue lysates extracted from growing (p25OE) vs. arrested (p25OFF) tumors. Polyclonal antibodies were used for probing the indicated phosphosites; mMTC = mouse MTC.

**Figure S6.**
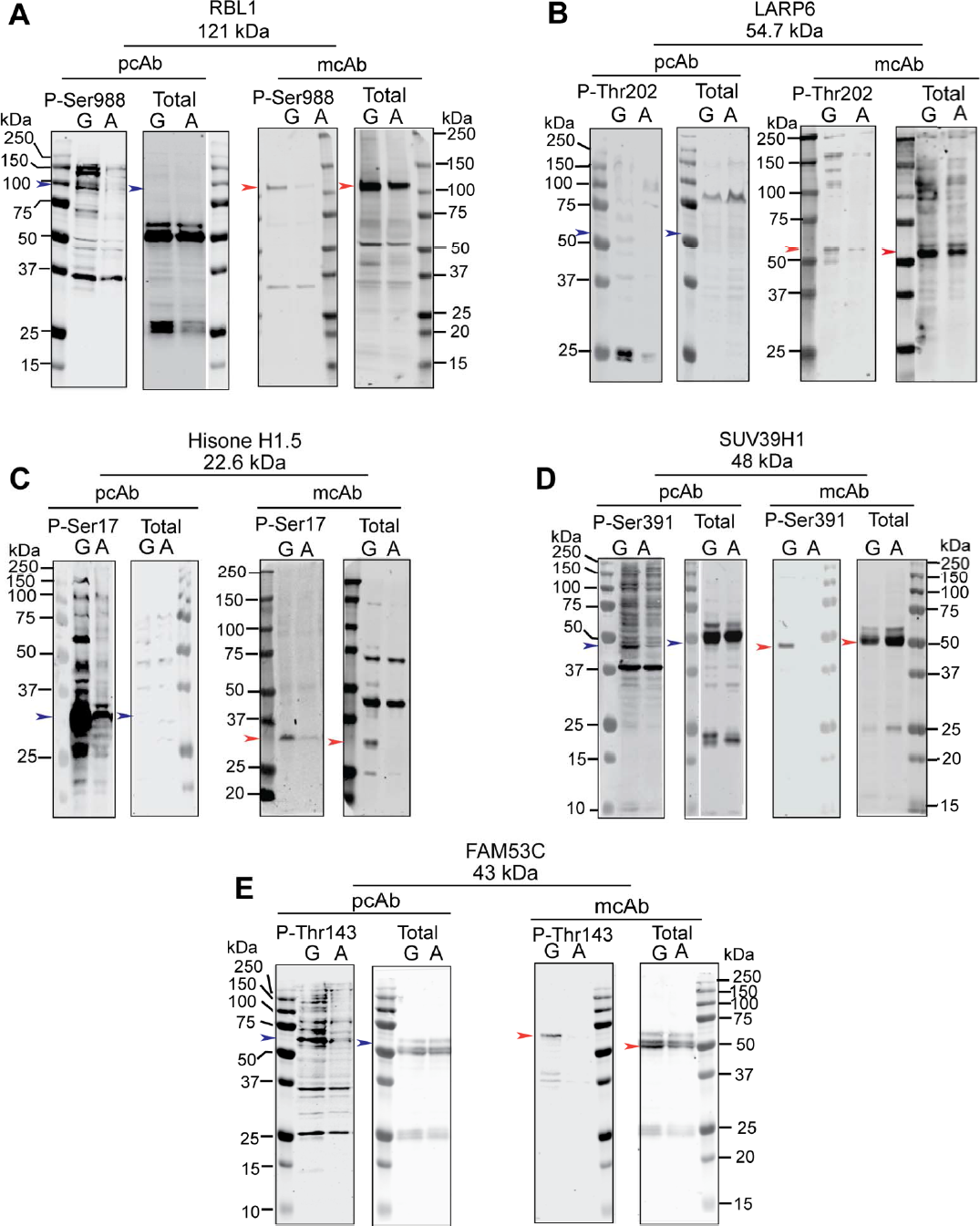
Characterization of recombinant phospho-specific mcAbs in mouse tumors. Comparative analysis of Cdk5 phosphorylation state-specific polyclonal antibodies (pcAb) vs. recombinant mcAbs. Lysates from growing (G) and arrested (A) tumors were analyzed via immunoblotting using Abs as indicated. Phospho-sites probed were (A) phospho-Ser988 RBL1, (B) phospho-Thr202 LARP6, (C) phospho-Ser17 Histone H1.5, (D) phospho-Ser391 SUV39H1, and (E) phospho-Thr143 FAM53C.

## Supplemental information

Download all supplemental files included with this article.

Table S1. Data summary of mutational profile of mouse tumors, related to Figure 2.

Table S2. Summary of overlapping mutated genes in mouse and human tumors, related to Figure 3.

Table S3. List of differentially expressed genes (DEGs) in NSE-p25OE and CGRP-p25OE mice, related to Figure 4.

Table S4 and S5. List of gene hits associated with the main clusters of upregulated DEGs in NSE-p25OE and CGRP-p25OE tumors respectively, related to Figure 4.

Table S6. List of altered genes that correlate with changes in mRNA expression of mouse tumors, related to Figure 5.

## References

1. Modigliani, E., et al., Prognostic factors for survival and for biochemical cure in medullary thyroid carcinoma: results in 899 patients. The GETC Study Group. Groupe d’etude des tumeurs a calcitonine. Clin Endocrinol (Oxf), 1998. 48(3): p. 265–73.

2. Bartz-Kurycki, M.A., O.E. Oluwo, and L.F. Morris-Wiseman, Medullary thyroid carcinoma: recent advances in identification, treatment, and prognosis. Ther Adv Endocrinol Metab, 2021. 12: p. 20420188211049611.

3. Maxwell, J.E., et al., Medical management of metastatic medullary thyroid cancer. Cancer, 2014. 120(21): p. 3287–301.

4. Sippel, R.S., M. Kunnimalaiyaan, and H. Chen, Current management of medullary thyroid cancer. Oncologist, 2008. 13(5): p. 539–47.

5. Chen, H., et al., The North American Neuroendocrine Tumor Society consensus guideline for the diagnosis and management of neuroendocrine tumors: pheochromocytoma, paraganglioma, and medullary thyroid cancer. Pancreas, 2010. 39(6): p. 775–83.

6. Pajak, C., et al., (68)Ga-DOTATATE-PET shows promise for diagnosis of recurrent or persistent medullary thyroid cancer: A systematic review. Am J Surg, 2022. 224(2): p. 670–675.

7. Chen, H., et al., Effective long-term palliation of symptomatic, incurable metastatic medullary thyroid cancer by operative resection. Ann Surg, 1998. 227(6): p. 887–95.

8. Oczko-Wojciechowska, M., et al., Current status of the prognostic molecular markers in medullary thyroid carcinoma. Endocr Connect, 2020. 9(12): p. R251–R263.

9. Pozo, K. and J.A. Bibb, The Emerging Role of Cdk5 in Cancer. Trends Cancer, 2016. 2(10): p. 606–618.

10. Pozo, K., et al., The role of Cdk5 in neuroendocrine thyroid cancer. Cancer Cell, 2013. 24(4): p. 499–511.

11. Carter, A.M., et al., Cdk5 drives formation of heterogeneous pancreatic neuroendocrine tumors. Oncogenesis, 2021. 10(12): p. 83.

12. Gupta, P., et al., Genetic impairment of succinate metabolism disrupts bioenergetic sensing in adrenal neuroendocrine cancer. Cell Rep, 2022. 40(7): p. 111218.

13. Meyer, D.A., et al., Striatal dysregulation of Cdk5 alters locomotor responses to cocaine, motor learning, and dendritic morphology. Proc Natl Acad Sci U S A, 2008. 105(47): p. 18561–6.

14. Lange, S., et al., Analysis pipelines for cancer genome sequencing in mice. Nat Protoc, 2020. 15(2): p. 266–315.

15. Bolger, A.M., M. Lohse, and B. Usadel, Trimmomatic: a flexible trimmer for Illumina sequence data. Bioinformatics, 2014. 30(15): p. 2114–20.

16. Li, H. and R. Durbin, Fast and accurate short read alignment with Burrows-Wheeler transform. Bioinformatics, 2009. 25(14): p. 1754–60.

17. McKenna, A., et al., The Genome Analysis Toolkit: a MapReduce framework for analyzing next-generation DNA sequencing data. Genome Res, 2010. 20(9): p. 1297–303.

18. Benjamin, D., et al., Calling Somatic SNVs and Indels with Mutect2. 2019, bioRxiv.

19. Kuilman, T., et al., CopywriteR: DNA copy number detection from off-target sequence data. Genome Biol, 2015. 16: p. 49.

20. Chen, S., et al., fastp: an ultra-fast all-in-one FASTQ preprocessor. Bioinformatics, 2018. 34(17): p. i884–i890.

21. Dobin, A., et al., STAR: ultrafast universal RNA-seq aligner. Bioinformatics, 2013. 29(1): p. 15–21.

22. Anders, S., P.T. Pyl, and W. Huber, HTSeq--a Python framework to work with high-throughput sequencing data. Bioinformatics, 2015. 31(2): p. 166–9.

23. Love, M.I., W. Huber, and S. Anders, Moderated estimation of fold change and dispersion for RNA-seq data with DESeq2. Genome Biol, 2014. 15(12): p. 550.

24. Zhuang, Y., et al., The novel function of tumor protein D54 in regulating pyruvate dehydrogenase and metformin cytotoxicity in breast cancer. Cancer Metab, 2019. 7: p. 1.

25. Baetscher, M., et al., SV40 T antigen transforms calcitonin cells of the thyroid but not CGRP-containing neurons in transgenic mice. Oncogene, 1991. 6(7): p. 1133–8.

26. Johnston, D., D. Hatzis, and M.E. Sunday, Expression of v-Ha-ras driven by the calcitonin/calcitonin gene-related peptide promoter: a novel transgenic murine model for medullary thyroid carcinoma. Oncogene, 1998. 16(2): p. 167–77.

27. Grundker, C. and G. Emons, The Role of Gonadotropin-Releasing Hormone in Cancer Cell Proliferation and Metastasis. Front Endocrinol (Lausanne), 2017. 8: p. 187.

28. Liu, H., et al., The oxytocin receptor signalling system and breast cancer: a critical review. Oncogene, 2020. 39(37): p. 5917–5932.

29. Stepulak, A., et al., Glutamate and its receptors in cancer. J Neural Transm (Vienna), 2014. 121(8): p. 933–44.

30. Haas, H.S., et al., The influence of glutamate receptors on proliferation and metabolic cell activity of neuroendocrine tumors. Anticancer Res, 2013. 33(4): p. 1267–72.

31. Wen, Y.A., et al., Downregulation of SREBP inhibits tumor growth and initiation by altering cellular metabolism in colon cancer. Cell Death Dis, 2018. 9(3): p. 265.

32. Wong, A., K. Nabata, and S.M. Wiseman, Medullary thyroid carcinoma: a narrative historical review. Expert Rev Anticancer Ther, 2022. 22(8): p. 823–834.

33. Minna, E., et al., Medullary Thyroid Carcinoma Mutational Spectrum Update and Signaling-Type Inference by Transcriptional Profiles: Literature Meta-Analysis and Study of Tumor Samples. Cancers (Basel), 2022. 14(8).

34. Cakir, M. and A.B. Grossman, Medullary thyroid cancer: molecular biology and novel molecular therapies. Neuroendocrinology, 2009. 90(4): p. 323–48.

35. Qu, N., et al., Genomic and Transcriptomic Characterization of Sporadic Medullary Thyroid Carcinoma. Thyroid, 2020. 30(7): p. 1025–1036.

36. Cerrato, A., V. De Falco, and M. Santoro, Molecular genetics of medullary thyroid carcinoma: the quest for novel therapeutic targets. J Mol Endocrinol, 2009. 43(4): p. 143–55.

37. Manfredi, G.I., et al., PI3K/Akt/mTOR signaling in medullary thyroid cancer: a promising molecular target for cancer therapy. Endocrine, 2015. 48(2): p. 363–70.

38. Yin, A., et al., Overexpression of NDRG2 Increases Iodine Uptake and Inhibits Thyroid Carcinoma Cell Growth In Situ and In Vivo. Oncol Res, 2016. 23(1-2): p. 43–51.

39. Karger, S., et al., FOXO3a: a novel player in thyroid carcinogenesis? Endocr Relat Cancer, 2009. 16(1): p. 189–99.

40. Cook, M., X.M. Yu, and H. Chen, Notch in the development of thyroid C-cells and the treatment of medullary thyroid cancer. Am J Transl Res, 2010. 2(1): p. 119–25.

41. Valenciaga, A., et al., Reduced Retinoblastoma Protein Expression Is Associated with Decreased Patient Survival in Medullary Thyroid Cancer. Thyroid, 2017. 27(12): p. 1523–1533.

42. Jajin, M.G., et al., Gas chromatography-mass spectrometry-based untargeted metabolomics reveals metabolic perturbations in medullary thyroid carcinoma. Sci Rep, 2022. 12(1): p. 8397.

43. Lukey, M.J., et al., The oncogenic transcription factor c-Jun regulates glutaminase expression and sensitizes cells to glutaminase-targeted therapy. Nat Commun, 2016. 7: p. 11321.

44. Chiefari, E., et al., Increased expression of AP2 and Sp1 transcription factors in human thyroid tumors: a role in NIS expression regulation? BMC Cancer, 2002. 2: p. 35.

45. Giuliani, C., I. Bucci, and G. Napolitano, The Role of the Transcription Factor Nuclear Factor-kappa B in Thyroid Autoimmunity and Cancer. Front Endocrinol (Lausanne), 2018. 9: p. 471.

46. Puig-Oliveras, A., et al., Expression-based GWAS identifies variants, gene interactions and key regulators affecting intramuscular fatty acid content and composition in porcine meat. Sci Rep, 2016. 6: p. 31803.

47. Motamed, M., et al., Steroid receptor coactivator 1 is an integrator of glucose and NAD+/NADH homeostasis. Mol Endocrinol, 2014. 28(3): p. 395–405.

48. Carter, A.M., et al., Phosphoprotein-based biomarkers as predictors for cancer therapy. Proc Natl Acad Sci U S A, 2020. 117(31): p. 18401–18411.

49. Lin, H., et al., Cdk5 regulates STAT3 activation and cell proliferation in medullary thyroid carcinoma cells. J Biol Chem, 2007. 282(5): p. 2776–84.

50. Haga, R.B. and A.J. Ridley, Rho GTPases: Regulation and roles in cancer cell biology. Small GTPases, 2016. 7(4): p. 207–221.

51. Rodriguez-Hernandez, I., et al., Rho, ROCK and actomyosin contractility in metastasis as drug targets. F1000Res, 2016. 5.

52. Heng, Y.W. and C.G. Koh, Actin cytoskeleton dynamics and the cell division cycle. Int J Biochem Cell Biol, 2010. 42(10): p. 1622–33.

53. Vasioukhin, V. and E. Fuchs, Actin dynamics and cell-cell adhesion in epithelia. Curr Opin Cell Biol, 2001. 13(1): p. 76–84.

54. Giardino, E., et al., Cofilin is a mediator of RET-promoted medullary thyroid carcinoma cell migration, invasion and proliferation. Mol Cell Endocrinol, 2019. 495: p. 110519.

55. Strock, C.J., et al., Cyclin-dependent kinase 5 activity controls cell motility and metastatic potential of prostate cancer cells. Cancer Res, 2006. 66(15): p. 7509–15.

56. Xie, Z., et al., Serine 732 phosphorylation of FAK by Cdk5 is important for microtubule organization, nuclear movement, and neuronal migration. Cell, 2003. 114(4): p. 469–82.

57. Pacifico, F. and A. Leonardi, Role of NF-kappaB in thyroid cancer. Mol Cell Endocrinol, 2010. 321(1): p. 29–35.

58. Currie, E., et al., Cellular fatty acid metabolism and cancer. Cell Metab, 2013. 18(2): p. 153–61.

59. Ferraro, G.B., et al., Fatty Acid Synthesis Is Required for Breast Cancer Brain Metastasis. Nat Cancer, 2021. 2(4): p. 414–428.

60. Do, P.A. and C.H. Lee, The Role of CDK5 in Tumours and Tumour Microenvironments. Cancers (Basel), 2020. 13(1).

61. Lowman, X.H., et al., The proapoptotic function of Noxa in human leukemia cells is regulated by the kinase Cdk5 and by glucose. Mol Cell, 2010. 40(5): p. 823–33.

62. Choi, J.H., et al., Anti-diabetic drugs inhibit obesity-linked phosphorylation of PPARgamma by Cdk5. Nature, 2010. 466(7305): p. 451-6.

63. Ciraku, L., et al., O-GlcNAc transferase regulates glioblastoma acetate metabolism via regulation of CDK5-dependent ACSS2 phosphorylation. Oncogene, 2022. 41(14): p. 2122–2136.

64. Dominguez, D., et al., A high-resolution transcriptome map of cell cycle reveals novel connections between periodic genes and cancer. Cell Res, 2016. 26(8): p. 946–62.

65. Pezzani, R., et al., Novel Prognostic Factors Associated with Cell Cycle Control in Sporadic Medullary Thyroid Cancer Patients. Int J Endocrinol, 2019. 2019: p. 9421079.

66. Valenciaga, A., et al., Transcriptional targeting of oncogene addiction in medullary thyroid cancer. JCI Insight, 2018. 3(16).

67. Yang, X., et al., TPX2 overexpression in medullary thyroid carcinoma mediates TT cell proliferation. Pathol Oncol Res, 2014. 20(3): p. 641–8.

68. Tuccilli, C., et al., Preclinical testing of selective Aurora kinase inhibitors on a medullary thyroid carcinoma-derived cell line. Endocrine, 2016. 52(2): p. 287–95.

69. Liu, M., C. Tolg, and E. Turley, Dissecting the Dual Nature of Hyaluronan in the Tumor Microenvironment. Front Immunol, 2019. 10: p. 947.

70. Slattery, M.L., et al., Genetic variation in RPS6KA1, RPS6KA2, RPS6KB1, RPS6KB2, and PDK1 and risk of colon or rectal cancer. Mutat Res, 2011. 706(1-2): p. 13-20.

71. Bignone, P.A., et al., RPS6KA2, a putative tumour suppressor gene at 6q27 in sporadic epithelial ovarian cancer. Oncogene, 2007. 26(5): p. 683–700.

72. Atas, E., M. Oberhuber, and L. Kenner, The Implications of PDK1-4 on Tumor Energy Metabolism, Aggressiveness and Therapy Resistance. Front Oncol, 2020. 10: p. 583217.

73. Han, H., et al., TRRUST: a reference database of human transcriptional regulatory interactions. Sci Rep, 2015. 5: p. 11432.

74. Zhou, Y., et al., Metascape provides a biologist-oriented resource for the analysis of systems-level datasets. Nat Commun, 2019. 10(1): p. 1523.

