## Supplementary material for "Faulty Metabolism: A Potential Instigator of an Aggressive Phenotype in Cdk5-dependent Medullary Thyroid Carcinoma": List of gene hits associated with the main clusters of upregulated DEGs in NSE-p25OE and CGRP-p25OE tumors respectively, related to Figure 4.

| Gene In GO And Hit List | Hits NSE-p25OE | Description | Category | LogP |
|---|---|---|---|---|
| 56 | ADAM8\|ARG1\|ARG2\|RHOH\|RUNX3\|CDKN1A\|CCN2\|EPHB2\|FCGR2B\|FCGR3A\|FN1\|GATA1\|HMOX1\|IGF1\|IGF2\|IL4 R\|INHBA\|ITGAM\|JUNB\|JUND \|LBP\|LGALS3\|MMP8\|PDGFB \|CCL3\|SHB\|SOX12\|TFRC\|THBS1\|TLR4\|TNFSF4\|TNFRSF 4\|ZAP70\|MAD1L1\|NCK2\|TNFSF14\|SPHK1\|BCL10\|TNFSF18\|SIRPB1\|MAD2L2\|MERTK\|SPINK5\|RIPK3\|CORO1A\|PAXIP1\|TNFRSF21\|IL20RB\|AKIRIN2\|CYP26B1\|CD177\|HAMP\|LOXL3\|DCAF15\|NLRP3\|TNFRSF13C | regulation of cell activation | GO Biological Processes | -12.77060957 |
| 55 | ADAM8\|ADRA2A\|BDKRB1\|CLDN4\|CSF2\|DEFB1\|EDN3\|EPHB2\|F3\|F7\|F10\|FOXC2\|FN 1\|FUT1\|HMOX1\|HRAS\|IGF1\|ITGA2\|ITGA5\|JUN\|LBP\|LGALS3\|MMP2\|MMP9\|SERPINE1\|PDGFB\|PFN1\|PIK3CD\|PTGS2\|RET\|CCL3\|CCL24\|SEMA3F\|THBS1\|TLR4\|VEGFA\|VEGFB\|WNT7A\|WNT11\|TNFSF14\|SPHK1\|TNFSF18\|SPOCK2\|CORO1A\|FERMT1\|PDGF C\|CAMK1D\|CEMIP\|S100A14\|ARHGEF39\|CARMIL2\|CCBE 1\|TAC4\|CXCL17\|SMIM22 | positive regulation of cell motility | GO Biological Processes | -13.97891985 |
| 52 | ADAM8\|ADRA2A\|BDKRB1\|CLDN4\|CSF2\|EDN3\|EPHB2\|F3\|F7\|F10\|FOXC2\|FN1\|FUT1\|HMOX1\|HRAS\|IGF1\|ITGA2\|ITGA5\|JUN\|LBP\|LGALS3\|MMP2\|MMP9\|SERPINE1\|PDGFB\|PFN1\|PIK3CD\|PTGS2\|RET\|CCL3\|CCL24\|SEMA3F\|THBS 1\|TLR4\|VEGFA\|VEGFB\|WNT7A\|WNT11\|TNFSF14\|SPHK1\|TNFSF18\|CORO1A\|FERMT1\|PDGFC\|CAMK1D\|CEMIP\|S100A14\|ARHGEF39\|CARMIL2\|CCBE1\|CXCL17\|SMIM22 | positive regulation of cell migration | GO Biological Processes | -13.01864966 |
| 42 | ADAM8\|ARG1\|ARG2\|RHOH\|RUNX3\|CDKN1A\|EPHB2\|FCGR2B\|FCGR3A\|IGF1\|IGF2\|IL4R\|INHBA\|JUNB\|LGALS3\|SHB\|SOX12\|TFRC\|TLR4\|TNFSF4\|TNFRSF4\|ZAP70\|MAD1L1\|NCK2\|TNFSF14\|BCL10\|TNFSF18\|SIRPB1\|MAD2L2\|MERTK\|SPINK5\|RIPK3\|CORO1A\|PAXIP1\|TNFRSF21\|IL20RB\|AKIRIN2\|CYP26B1\|LOXL 3\|DCAF15\|NLRP3\|TNFRSF1 3C | regulation of lymphocyte activation | GO Biological Processes | -9.402397222 |
| 40 | ADAM8\|APOA1\|RHOH\|RUNX 3\|DMP1\|FOXC2\|FN1\|FUT1\|GCNT1\|IGF1\|IGF2\|IL4R\|ITGA2\|ITGA5\|LIF\|MIP\|P4HB\|PDGFB\|PLAUR\|SERPINF2\|RET \|SAA1\|SHB\|SOX12\|TFRC\|TNFSF4\|VEGFA\|ZAP70\|NCK2\|ADAM19\|TNFSF14\|BCL10\|TNFSF18\|SPOCK2\|SIRPB1\|CORO1A\|KIF26B\|FERMT1\|NLRP3\|TNFRSF13C | positive regulation of cell adhesion | GO Biological Processes | -8.906493555 |
| 40 | ADAM8\|APOA1\|ARG1\|ARG2\|RHOH\|ASS1\|RUNX3\|FCGR2 B\|GCNT1\|IGF1\|IGF2\|IL4R\|LGALS3\|PCDH8\|PLAUR\|SERPINF2\|SHB\|SOX12\|TFRC\|TNFSF4\|VEGFA\|ZAP70\|MAD1L1\|NCK2\|ADAM19\|TNFSF14\|BCL10\|TNFSF18\|SIRPB1\|MAD2L2\|CORO1A\|TNFRSF21\|FXYD5\|IL20RB\|KIF26B\|AKNA\|LOXL3\|NLRP3\|TNFRSF13 C\|MDGA1 | regulation of cell-cell adhesion | GO Biological Processes | -8.675154491 |
| 35 | ADAM8\|RHOH\|RUNX3\|CDKN1A\|EPHB2\|FCGR3A\|GATA 1\|IGF1\|IGF2\|IL4R\|ITGAM\|JU ND\|LBP\|MMP8\|CCL3\|SHB\|SOX12\|TFRC\|THBS1\|TLR4\|TNFSF4\|TNFRSF4\|ZAP70\|NCK2\|TNFSF14\|BCL10\|SIRPB1 \|MAD2L2\|CORO1A\|PAXIP1\|AKIRIN2\|CD177\|HAMP\|NLR P3\|TNFRSF13C | positive regulation of leukocyte activation | GO Biological Processes | -9.138742901 |
| 31 | ADAM8\|ARG1\|ARG2\|RHOH\|RUNX3\|FCGR2B\|IGF1\|IGF2\|IL4R\|JUNB\|LGALS3\|SHB\|SOX12\|TFRC\|TNFSF4\|ZAP70\|MAD1L1\|NCK2\|TNFSF14\|BCL10\|TNFSF18\|SIRPB1\|SPINK5\|RIPK3\|CORO1A\|TNFRSF21\|IL20RB\|CYP26B1\|LOXL3\|NLRP3\|TNFRSF13C | regulation of T cell activation | GO Biological Processes | -6.956152205 |
| 29 | ADAM8\|ARG1\|ARG2\|RHOH\|ASS1\|RUNX3\|FCGR2B\|GCNT1\|IGF1\|IGF2\|IL4R\|LGALS3\|SHB\|SOX12\|TFRC\|TNFSF4\|ZAP70\|MAD1L1\|NCK2\|TNFSF14\|BCL10\|TNFSF18\|SIRPB1\|CORO1A\|TNFRSF21\|IL20RB\|LOXL3\|NLRP3\|TNFRSF13C | regulation of leukocyte cell-cell adhesion | GO Biological Processes | -5.930947343 |
| 22 | ARG1\|ARG2\|CDKN1A\|EPHB 2\|FCGR2B\|FCGR3A\|IGF1\|IGF2\|LGALS3\|TFRC\|TLR4\|TNFSF4\|TNFRSF4\|ZAP70\|MAD1L1\|NCK2\|TNFSF18\|RIPK3\|CORO1A\|TNFRSF21\|IL20RB \|TNFRSF13C | regulation of lymphocyte proliferation | GO Biological Processes | -5.923735243 |
| 7 | CSF3\|PFN1\|CCL24\|NCK2\|BAIAP2\|BAIAP2L2\|CARMIL2 | positive regulation of actin filament polymerization | GO Biological Processes | -3.391841492 |
