## Supplementary material for "Faulty Metabolism: A Potential Instigator of an Aggressive Phenotype in Cdk5-dependent Medullary Thyroid Carcinoma": List of gene hits associated with the main clusters of upregulated DEGs in NSE-p25OE and CGRP-p25OE tumors respectively, related to Figure 4.

| Gene In GO And Hit List | Hits CGRP-p25OE | Description | Category | LogP |
|---|---|---|---|---|
| 40 | CETN2\|HSP90AA1\|POLR2K\|PPP1CC\|PRKAR2B\|PSMA1\|PSMC6\|RB1\|RBBP7\|RFC1\|NSD2\|SEM1\|SMC1A\|H2AC14\|H2BC15\|H2BC21\|H4C1\|H4C11\|H3C7\|SMC3\|CEP57\|CKAP5\|POLD3\|ARPP19\|NUP160\|ITGB3BP\|SUN2\|GMNN\|FZR1\|NUP54\|NDE1\|CHTF8\|CEP192\|HAUS2\|CENPJ\|POLE4\|MIS12\|CEP70\|CEP41\|MZT1 | Cell Cycle | Reactome Gene Sets | -10.73571513 |
| 37 | CETN2\|FMR1\|HSP90AA1\|MSH2\|NAP1L1\|NONO\|PMS1\|PMS2\|RBBP7\|RFC1\|SSRP1\|TP73\|NSD2\|SMC1A\|TTF2\|SMC3\|POLD3\|PTGES3\|GMNN\|FZR1\|CHTF8\|HPF1\|FANCL\|NUDT15\|SMPD3\|POLE4\|INIP\|SMC6\|H2AC25\|PIWIL4\|COMMD1\|MCMDC2\|USP51\|APLF\|PRIMPOL\|MCM9\|MEI4 | DNA metabolic process | GO Biological Processes | -7.797726637 |
| 25 | FMR1\|HNRNPA1\|HNRNPC\|NONO\|EXOSC10\|SRSF3\|SNRPD3\|TAF9\|TAF13\|TTF2\|GEMIN2\|PRPF4\|PTBP3\|SAP18\|SYNCRIP\|KHDRBS3\|KHDRBS1\|SF3A3\|PSIP1\|DIS3\|USP49\|SNIP1\|THOC7\|HNRNPLL\|NT5C3B | mRNA metabolic process | GO Biological Processes | -4.14937088 |
| 24 | ATP5PB\|ATP5ME\|ATP5PO\|DCTD\|GNAI3\|GUCY2C\|HMGCS1\|HTR2A\|IMPDH2\|NDUFS4\|NDUFS5\|PRPS2\|SDHD\|ELOVL4\|NME5\|KCNAB2\|ATP5MJ\|PAICS\|ACSL5\|CMPK1\|ELOVL2\|NUDT15\|NT5C3B\|CMPK2 | nucleotide metabolic process | GO Biological Processes | -5.037707401 |
| 21 | HDAC2\|RBBP7\|SMARCC1\|SMARCD2\|TAF9\|NSD2\|H2AC15\|H2AC14\|H2AC16\|H2BC15\|H2BC21\|H4C1\|H4C11\|H3C7\|SAP18\|MSL3\|ELP6\|PRDM16\|MEAF6\|DPY30\|H2AC25 | Chromatin organization | Reactome Gene Sets | -7.963690505 |
| 16 | FBN2\|GATA4\|HDAC2\|HIF1A\|RBBP7\|NREP\|ONECUT2\|SULF1\|ASPN\|SINHCAF\|PRDM16\|ADAMTS12\|CD109\|BMPER\|HTRA4\|ELAPOR2 | regulation of cellular response to growth factor stimulus | GO Biological Processes | -3.5317087 |
| 12 | CETN2\|HDAC2\|HNRNPC\|SP3\|SMC1A\|H4C1\|H4C11\|SMC3\|CASP8AP2\|NUP160\|NUP54\|SMC6 | SUMO E3 ligases SUMOylate target proteins | Reactome Gene Sets | -4.168744437 |
| 11 | CCNG1\|CSE1L\|HDAC2\|MSH2\|PMS2\|RB1\|SP1\|TAF9\|TP73\|PYCARD\|SCN3B | PID P53 DOWNSTREAM PATHWAY | Canonical Pathways | -4.647902875 |
| 11 | NAP1L1\|RBBP7\|RFC1\|SSRP1\|POLD3\|GMNN\|CHTF8\|POLE4\|MCMDC2\|PRIMPOL\|MCM9 | DNA replication | GO Biological Processes | -3.267596633 |
| 10 | FBN2\|HDAC2\|RBBP7\|NREP\|ONECUT2\|ASPN\|SINHCAF\|PRDM16\|CD109\|HTRA4 | regulation of transforming growth factor beta receptor signaling pathway | GO Biological Processes | -3.654567892 |
| 6 | CDC42\|FMR1\|RALA\|FNBP1L\|TENM2\|DOCK11 | regulation of filopodium assembly | GO Biological Processes | -3.670185807 |
| 4 | CAPZA1\|CDC42\|RALA\|TCP1 | Gene and protein expression by JAK-STAT signaling after Interleukin-12 stimulation | Reactome Gene Sets | -2.423162463 |
